# Nuclear deformation and dynamics of migrating cells in 3D confinement reveal adaptation of pulling and pushing forces

**DOI:** 10.1101/2023.10.30.564765

**Authors:** Stefan Stöberl, Johannes Flommersfeld, Maximilian M. Kreft, Martin Benoit, Chase P. Broedersz, Joachim O. Rädler

## Abstract

Eukaryotic cells show an astounding ability to migrate through pores and constrictions smaller than their nuclear diameter. However, the forces engaged in nuclear deformation and their effect on confined cell dynamics remain unclear. Here, we study the mechanics and dynamics of nuclei of mesenchymal cancer cells as they spontaneously and repeatedly transition through 3D compliant hydrogel channels. We find a biphasic dependence of migration speed and transition frequency on channel width, revealing maximal transition rates at widths comparable to the nuclear diameter. Using confocal imaging and hydrogel bead displacement, we determine the nuclear deformation and corresponding forces during spontaneous confined migration. We find the nucleus to reversibly deform with an elastic modulus not adapting to the confinement. Instead, with decreasing channel width, the nuclear shape during transmigration changes biphasically concomitant with the transitioning dynamics. The nucleus exhibits a prolate form along the migration direction in wide channels and a more compressed oblate shape in narrow channels. We propose a physical model for confined cell migration that explains the observed nuclear shapes and slowing down in terms of the cytoskeletal force-generation adapting from a pulling-to a pushing-dominated mechanism with increasing nuclear confinement.

## Main

Cell migration plays a key role in physiological processes such as wound healing, immune response, and cancer metastasis^1–4^. Cancer cell invasion is an essential step in the metastatic process, owing to the complex interplay of the molecular control of protrusions, reorganization of the cytoskeleton and interaction with the extracellular matrix (ECM). As cells invade this matrix, they squeeze through tight meshwork forming confinements ranging from less than 1 µm up to tens of microns^5,6^. By invading constrictions smaller than the nuclear diameter, the nucleus is required to deform to the size of the constriction^7–9^. The need of physical deformation emphasizes the importance of nuclear flexibility in cellular invasion into the ECM network^7,10^. Migration in confining environments raises the general question, whether and how cells can control their nuclear properties and force generation machinery to overcome such varying spatial constrictions.

Over the past two decades, an increasing number of studies have focused on the role of confinement in cell migration. In general, restriction of two-dimensional migration to one-dimensional patterned lanes or microstructured channels was found to accelerate migration^11,12^. However, in narrow 3D constrictions where nuclear deformation is required, the translocation of the nucleus was identified as the rate-limiting step during migration in early studies^13–16^. Initially, the role of the nucleus in confined migration was reduced to that of a passive cargo, whereby its viscoelastic properties were important for cellular migration^7,17,18^. Recent studies using flat silicon microcantilevers to confine cells, however, have presented evidence that besides passively deforming, cells can use their nucleus to sense confinement and actively change nuclear shape^19,20^. In contrast to the externally imposed confinement in these studies, cells typically ‘self-impose’ their confinement when spontaneously migrating within and through tight pores. It is still unclear whether such active mechanisms play a role during self-imposed confined migration.

To study self-imposed confined cell migration quantitatively, artificial microfluidic assays are commonly employed. Notably, microfabricated hydrogel-based migration assays^18,21–23^ and trans-well migration assays^17,24,25^ have proven valuable to understand the effect of nuclear mechanical properties on cell migration and survival at the single cell level^17^. Long PDMS channels^12,14,26,27^ are frequently utilized to investigate the migration dynamics of cells physically confined within a controlled experimental environment. However, to investigate cell entry into constrictions, shorter PDMS channels have proven instrumental, as they allow to study the effect of confinement on migration without perturbing the overall mode of migration^28^.

While the importance of the nucleus in confined migration has become clear, it remains debated how the cytoskeleton is involved in deforming and maneuvering the nucleus into constrictions. Several studies identified a dominant contribution of contractile actomyosin fibers in front of the nucleus generating “pulling” forces that are transmitted to the nucleus via the LINC complex anchoring actin via nesprin-2 to the nuclear membrane^29–31^. In contrast, other work highlighted the significance of rear cortical contractions during transmigration through narrow constrictions, representing a mechanism based on an osmotic “pushing” force^15,32,33^. These cortical contractions are thought to be triggered by strong nuclear deformations that lead to Ca^2+^ release, resulting in increased myosin recruitment to the cortex^19,20^. Together, these studies suggest an interplay between geometrical confinement, the cellular migration machinery, and nuclear mechanics. However, it is unclear how this interplay determines migration dynamics when cells self-impose physical confinement by spontaneously entering tight constrictions^7,19,20,34^.

Here, we introduce arrays of compliant 3D hydrogel micro-cavities to monitor single cancer cells spontaneously migrating through a constricting channel. We determine the nuclear dynamics, volume, deformations, and confinement forces for cells that repetitively migrate through a narrow channel, with constriction widths varying from the cell size to well below the nuclear diameter. For the cancerous cell line (MDA-MB-231) used in this study, we observe a bi-phasic behavior of cell transition rates as a function of constriction width with rates peaking around a channel width comparable to the nuclear diameter. Analyzing the full 3D nuclear deformation and forces during transitions, we find that the nucleus can be described as a compressible elastic object. Together, our results can be captured by a model in which nuclear deformation is determined by an adaptive force generation mechanism when cells encounter confinement. This mechanistic model explains the observed dependence of both the cellular migration dynamics and nuclear morphology on channel width. Our findings demonstrate that cells actively adapt their cytoskeleton force-generation also during ‘self-imposed’ cell migration to enable the nucleus to overcome a broad range of spatial constrictions.

## RESULTS

### Versatile hydrogel-based 3D confined migration assay

To quantitatively study the influence of physical confinement and nuclear deformations on the migration dynamics of cells, we develop a hydrogel-based migration assay to perform high-throughput cell migration experiments within a controllable geometry (Fig. 1a). Our setup uses 3D dumbbell-shaped micro-cavities embedded within a 20 µm thick PEG-NB hydrogel layer. Fluorescent nano-beads mixed into the hydrogel serve as markers to determine forces exerted by cells from the elastic deformation field of the walls (Fig. 1b). The bottom of the cavities is coated with fibronectin to promote cell adhesion, while the PEG-NB hydrogel is non-adhesive to cells (Fig. S1). This design ensures that cells migrate at the bottom of the cavity, while remaining entirely confined within the cavity walls throughout the experiments (Fig. 1b (ii)). Accordingly, the assay is a 3D and deformable generalization of the flat microcontact printed dumbbell pattern used in prior work^11^. We employ time-lapse phase-contrast microscopy to investigate repeated migration of cells from one cavity site to the other through the deformable channel with defined width. Metastatic human breast cancer cells (MDA-MB-231) were placed inside the 3D dumbbells, which we observe to migrate spontaneously in mesenchymal mode inside the micro-cavities (Fig. 1c). At the entrance of the constricting channel, lamellipodia-like protrusions develop repeatedly, occasionally evolving into larger sustained protrusions. While some protrusions retract rapidly, others reach the adjacent unoccupied cavity. In the new cavity the protrusion widens with an almost half-circular shape, which ultimately results in the transition of the cell body. After the complete cellular transition, the shape and actin distribution of the cytoskeleton randomizes (see Video S1), leading to a state identical to that preceding the transition. Subsequently, another transition soon follows in the reverse direction, illustrating the reversible nature of this confined migration (see Video S2 & S3).

**Figure 1:**
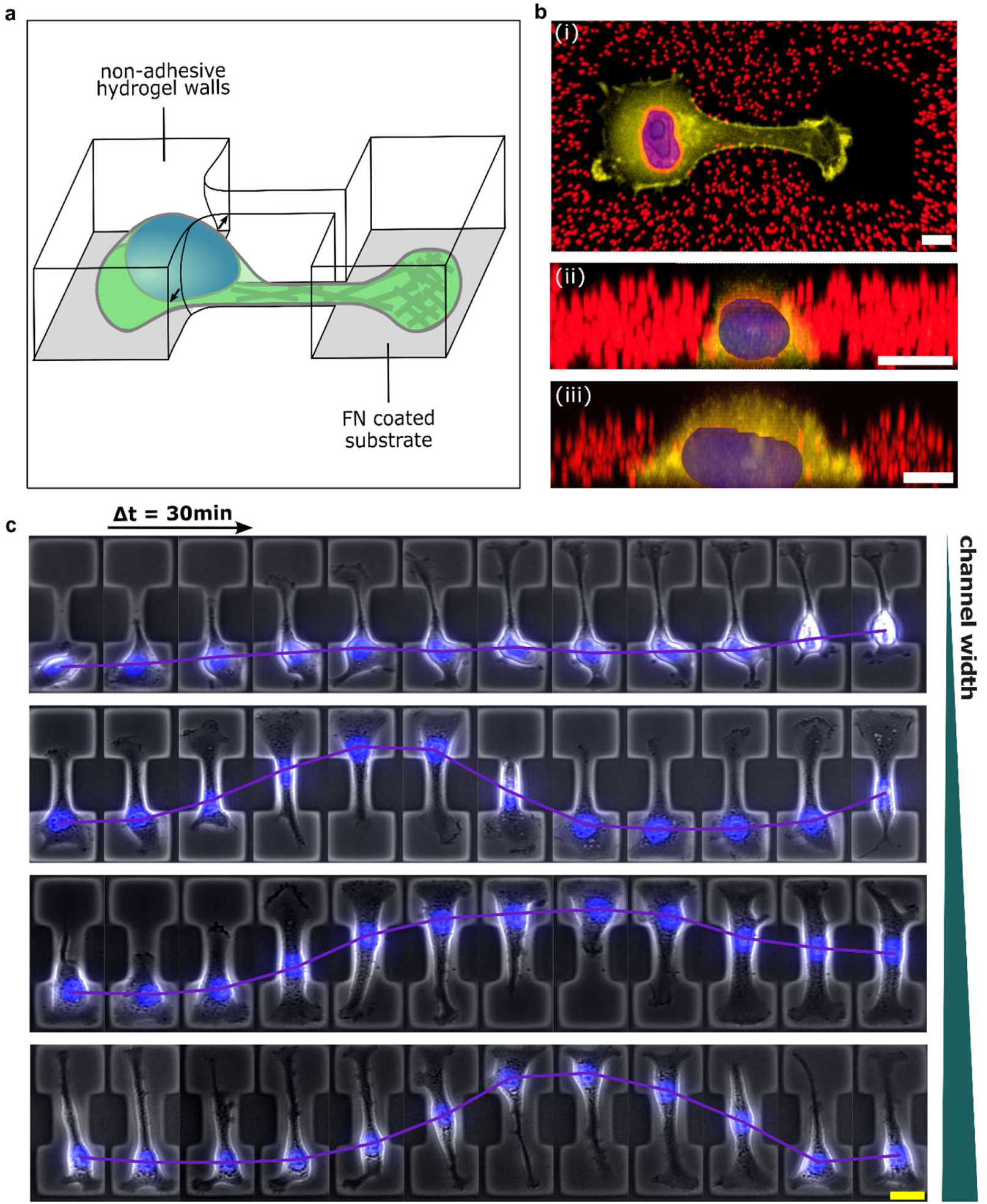
Synthetic hydrogel-based assay to study 3D confined cell migration. **(a)** Schematic depicting of the experimental assay consisting of two fibronectin-coated square chambers (35x35 μm²) connected by a channel. The dumbbell-shaped geometry is surrounded by non-adhesive and soft PEG-NB hydrogel, which gets deformed by cells traversing the constriction. **(b)** Exemplary confocal images show an MDA-MB-231 cell co-expressing fluorescently labeled histones (mcherry-H2B) and F-actin (LifeAct-TagGFP2) confined to the micropattern (i). Embedded fluorescent nano-beads indicate the surrounding PEG-NB hydrogel, which physically confines cells within the cavity (ii). As the hydrogel is non-adhesive, cell adhesion is solely restricted to the substrate (ii) (scale bar, 10 um). **(c)** Exemplary timeseries of MDA-MB-231 cells for varying constriction width (D = 3 – 35 um) (scale bar, 20 um).

### Cells show bi-phasic migration rates depending on confinement

We investigate cell migration as a function of confinement widths by fabricating 3D dumbbell arrays with channel width varying over a broad range from 4 to 35 µm. For channel widths wider than the nuclear diameter (Fig 1c, lowest row), cells can enter the channel without requiring nuclear deformations. In this case the migration within the channel shows little persistence and some cells even change their direction of motion in the middle of the channel without reaching the opposite side of the pattern (see Video S2). In channels with intermediate widths smaller than the nuclear diameter (Fig 1c, 2^nd^ and 3^rd^ row), cells exhibit large nuclear deformation (see Video S3). After nuclear shape adaption almost all cells consistently migrate to the other chamber. For the narrowest channels (Fig 1c, upper row), unsuccessful attempts of nuclear translocations are frequently observed. In channels where the nucleus becomes confined (<12µm), we observe an increased fraction of ‘trapped’ cells that typically still form protrusions that extend to the other side of the pattern but are unable to move their nucleus into the channel (Fig. 1c, upper row, Fig 2a, columns 1 and 2). Our results do not indicate a specific threshold width below which migration is completely inhibited, but rather a gradual reduction in the fraction of migratory cells with diminishing channel width. Moreover, when transitions require large nuclear deformations, the migration behavior markedly differ from dynamics found on flat micropatterned 2D dumbbells, where we previously observed frequent transitions, even for the narrowest bridge widths^35^. This observation highlights that 3D confinement qualitatively alters cell translocation.

**Figure 2:**
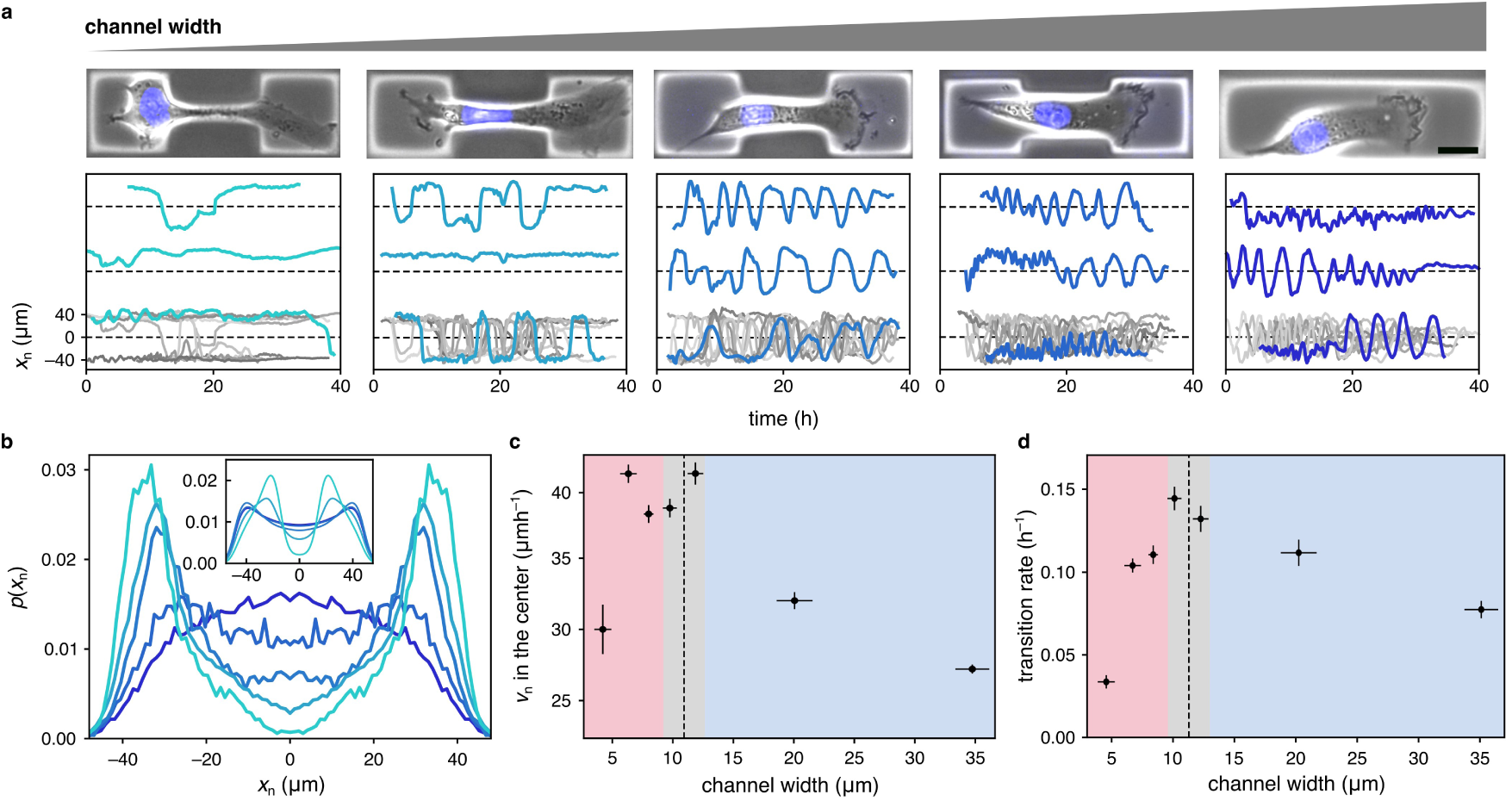
Migration statistics for varying channel widths. **(a)** Representative examples of cell trajectories for different channel widths (from left to right: 4 µm, 7 µm, 12 µm, 20 µm and 35 µm). **(b)** The distributions of the nuclear position *x_n_* at different channel widths. Same color scheme as in (a). **(c)** Average velocity in the center of the pattern (|*x_n_*|<=6µm). The maximal velocities are observed at an intermediate channel width comparable to the nuclear width. **(d)** Transition rates for varying channel widths. Similar to the nuclear velocities, the highest transition rates are observed at intermediate channel widths. Error bars in (b) and (c): error bars associated with the x-axis represent the standard deviation, while error bars associated with the y-axis represent the standard error.

To further investigate this, we evaluate the probability distribution of nuclear positions in our micro-cavities, the average velocity at the channel center, and the transition rates across varying channel widths. In the absence of confinement, the distribution of nuclear positions along the long axis of the pattern is nearly uniform over large parts of the micro-cavity (Fig. 2b). With decreasing channel width, the distribution transforms into a double peaked distribution, with marked maxima in the chambers and a pronounced local minimum located in the center of the confining channel. However, even for channels widths of 4 µm, substantially smaller than the average nuclear diameter, we still observe cells that can successfully move their nucleus through these remarkably tight constrictions. Interestingly, we observe a nonmonotonic dependence of the average velocity on the channel width, with a maximum at a confinement width comparable to the undeformed diameter of the nucleus (Fig. 2c). The overall transition kinetics for cells to enter and transmigrate the constricting channel, quantified by the transition rate, shows a similar biphasic dependence on channel width (Fig. 2d): When starting from the unconfined case and gradually decreasing channel width, we initially observe an increase of the transition rate, followed by a steep drop in the transition rate at channel widths below the nuclear size. We term these the ‘confinement-enhanced migration’ (CEM) and the ‘confinement-reduced migration (CRM) regimes. These results illustrate the effect that confinement has on the migration behavior of cells, where it either appears to enhance the rate of transitions at wider channel widths or strongly impede transitions when cells need to physically deform their nuclei in tight constrictions, as observed in other studies^15,36^.

### Nucleus deformation affects migrations dynamics

To obtain a better understanding of the underlying dynamics driving the transition from the CEM- to the CRM-regime, we exploit the statistics of nuclear velocities and transition rates obtained from our high throughput experimental assay. We use the recorded nuclear trajectories to infer a stochastic dynamical model for this confined migration process, enabling us to disentangle deterministic and stochastic components of the dynamics. Both, free and confined 2D migration can be described by an underdamped Langevin equation of the form ^11,37–39^

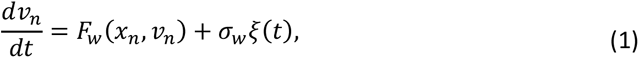

where *F_w_* (*x_n_*, *v_n_*) captures how the deterministic acceleration of the nucleus depends on position and velocity in a confinement of width *w*, and the Gaussian white noise *ξ*(*t*) of strength *σ_w_* accounts for the stochastic nature of cell migration. Here, we extend this approach to 3D confined migration by applying the Underdamped Langevin Inference algorithm^38^ on our data (SI section 3).

To understand the impact of the 3D confinement on dynamics, we compare the deterministic terms *F_w_* (*x_n_*, *v_n_*) inferred for different channel widths *w* (Fig. 3a). We focus on the transition from the CEM- into the CRM-regime, such that we only consider the dynamics at channel widths of 12 µm and below. For *w* ≥ 7 µm, we find qualitatively similar models: The nucleus on average strongly decelerates in the center of the two chambers. By contrast, as the nucleus approaches the constricting channel, it accelerates into the channel and ultimately transitions to the other side of the pattern. The structure of these nonlinear dynamics is at first glance reminiscent of cells migrating on corresponding 2D micro-patterns^11^. Interestingly, at the narrowest channel width (4 µm), this acceleration region vanishes. consistent with the more stationary behavior that we observed experimentally at this channel width (Fig. 2a).

**Figure 3:**
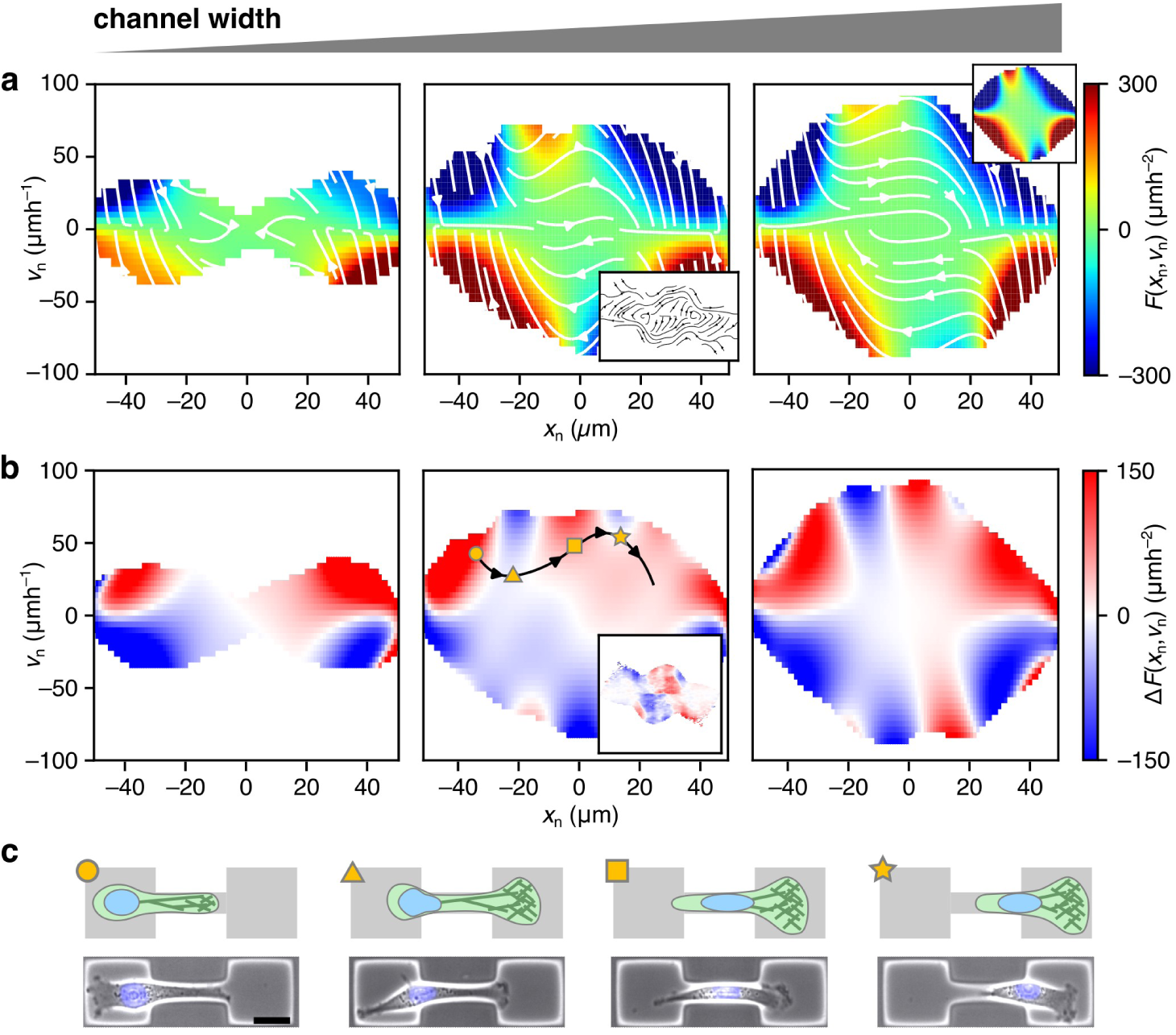
Inferred non-linear dynamics of the nucleus for varying channel widths (from left to right: 4 µm, 7 µm, 9 µm). **(a)** The inferred deterministic term *F_w_*(*x_n_*, *v_n_*) within the experimentally sampled region (see SI for details). The unsampled region is shown in white. Central inset: Predicted deterministic dynamics obtained from simulations of the mechanistic model (*w* = 7 µm). Right inset: *F_w_* (*x_n_*, *v_n_*) for a channel width of 12 µm. **(b)** The difference between the deterministic term *F_w_* (*x_n_*, *v_n_*) and the reference term at 12µm (‘Nuclear Confinement Maps’). The black line indicates an exemplary trajectory of a transitioning cell. Inset: Predicted confinement signature map obtained from simulations of the mechanist model (7 µm). **(c)** Snapshots of the typical cellular morphology at different points during the transition as indicated in (b) (scale bar: 20 µm).

To better visualize the effect of nuclear deformations on the non-linear dynamics, we compute the difference Δ*F_w_* = *F*_12µm_ − *F_w_* (Fig. 3b) to compare to the channel width *w* = 12 µm below which nuclear deformations become significant, which we term ‘Nuclear Confinement Maps’ (NCM). At all three considered channel widths, the NCMs show a number of shared qualitative features: A cell that migrates along a typical trajectory from the left to the right chamber (black line in Fig. 3b, center) starts in a region where Δ*F_w_* > 0 (circle in Fig. 3b, c). In previous work, we found that on 2D micro-patterns increasing confinement of the protrusion stimulates increasing protrusion growth and thus stronger accelerations of the nucleus towards the channel^35^. This pronounced region of Δ*F_w_* > 0 could be a signature of this ‘geometry adaptation’ mechanism also being present in 3D confinement. Once the nucleus approaches the channel, it crosses over into a region of Δ*F_w_* < 0 in the NCM (triangle in Fig. 3b, c). Since the region of Δ*F_w_* < 0 coincides with the point at which further migration requires large nuclear deformations, this suggest that this feature of the NCM may be due to an effective deformation energy barrier that impedes entry into the channel. Once the nucleus is moved into the channel, we again observe a region of acceleration (square in Fig. 3b, c) that is consistent with an elastic release of the tension that was build up to deform the nucleus in the previous step. If cells reach high enough velocities, they cross through another region of Δ*F_w_* < 0 (star in Fig. 3b, c) as it leaves the channel. Overall, the NCMs indicate that there is a clear qualitative signature of 3D nucleus confinement that suggest that elastic deformations of the nucleus affect the transmigration dynamics.

### Nuclei reversibly reduce volume during transmigration

To further investigate the role of nuclear mechanics indicated by the NCMs, we further analyze the nuclear deformations induced by the confining channel. We find that during transmigration the nuclear shape is strongly deformed depending on the channel width. Interestingly, the nucleus rapidly recovers to its original round shape, with a relaxation time of 20 minutes after exiting from the channel (Fig. S8). This observation suggests that in our set-up with relatively short time scales of deformation the nuclear response is predominantly elastic, consistent with the dynamics revealed by the NCMs. This is consistent with relatively low lamin A levels found in MDA-MB-231 cells^40^, which was shown to increase the reversibility of nuclear deformations^17^.

To quantify the full 3D nuclear deformation during transition events, we employ confocal microscopy (Fig. 4a, Fig. S2 and Video S4. Using rendering software, we determine the nuclear volume and ellipsoidal main axis (see methods). On longer time scales, over the duration of the experiments, we observe a continuous increase of the nuclear volume of both relaxed and compressed states. This trend corresponds to cell growth at a rate 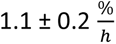 (Fig. S2a). As a control the nuclear volume growth in cavities without any constriction was measured (Fig. S2b) yielding a similar nuclear growth rate of 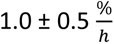. On shorter timescales, we observe a temporary reduction in nuclear volume, when the nucleus enters the channel, which repeats in subsequent transitions (Fig. 4a). From the full 3D deformation under confinement, we determine a Poisson ratio for MDA-MB-231 cell nuclei of *v_n_*=0.40 ± 0.02 independent of the channel width (Fig. 4a inset, SI section 1.6). Together, these results show that in the process of translocation the nucleus behaves as a compressible elastic material with slightly anisotropic response indicated by a Poisson ratio distinct from 0.5.

**Figure 4:**
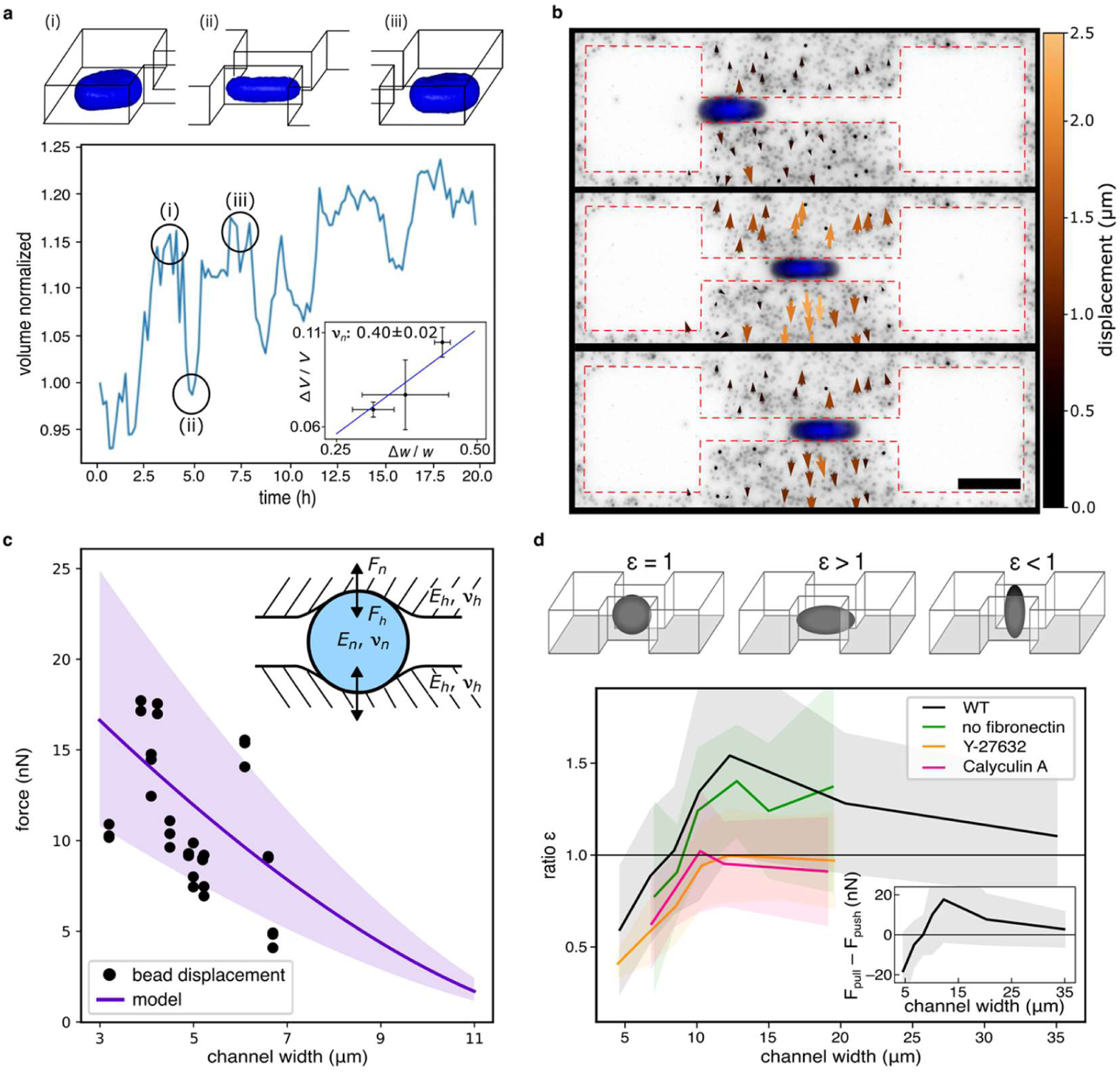
Mechanical properties and deformations of cell nuclei in confinement. **(a)** Quantitative nuclear volume analysis of exemplary confocal z-stacks over 20h. Nuclear volume decreases in confinement (ii) compared to an unconfined state (i and iii). Data were obtained using the software ‘Arivis4D’. Inset plot: Relative volume change of the nucleus when confined by the channel with three different widths (mean ± SD for n > 2 cell nuclei per channel width) and fit of an elastic model to obtain the Poisson ratio ν of the nucleus (blue line). **(b)** Representative image of the fluorescently labeled nucleus of an MDA-MB-231 cell passing through the soft PEG-NB hydrogel channel. Color-coded arrows indicate the displacement field of fluorescent beads embedded in the hydrogel (scale bar, 20 μm). **(c)** Forces exerted by the nucleus to the hydrogel walls as a function of channel width. For comparison, the expected forces are shown based on a Hertz model using independently measured Young’s moduli of the hydrogel and cell nucleus (shaded area indicates one standard deviation). Inlet shows schematics of the underlying Hertz model. **(d)** Relative change in the nuclear aspect ratio indicates an elongated (ε > 1) or compressed (ε < 1) nuclear shape in the direction of migration, which is consistent with pulling or pushing dominated translocation, respectively. Inset: difference between pulling and pushing forces acting on the nucleus. The shaded areas indicate one standard deviation.

### Nuclear elastic properties are independent of confinement

To further investigate the nuclear elastic properties and whether they are affected under strong confinement, we measure the forces exerted onto the channel walls by transitioning cells. We analyze the indentation of the soft hydrogel walls during transitions tracking the displacement of fluorescent beads embedded in the hydrogel (see Video S5). The observed bead displacements are decomposed into contributions perpendicular and parallel to the migration direction (Fig. S4). The normal and tangential forces are estimated by using the maximum bead displacement in the PEG-NB hydrogel whose Young’s modulus of 2.9 kPa we independently determined using Atomic Force Microscopy (AFM) with bead tips (Fig. S3). Within our detection limit, we did not observe tangential components, which is in agreement with the non-adhesive property of the PEGylated hydrogel. For the normal component we find a non-linear increase of the normal forces exerted by the nucleus with increasing confinement (Fig. 4c). For comparison, we include a theoretical prediction for the contact forces based on an adapted Hertz model with a soft wall indented by a deformable nucleus represented by an elastic sphere with Poisson’s ratio *v_n_*=0.40 and Young’s modulus *E_n_ =* 0.4 kPa, independently measured by AFM in the absence of the confinement (Fig. S3). The agreement between the estimated contact forces exerted on the nucleus and the predictions of the adapted Hertz model, which depends on the nucleus Young’s modulus that was determined in the absence of confinement, indicate that the elasticity properties of the nucleus do not show noticeable changes during the transmigration of a cell through a 3D constriction.

### Nuclear deformation reveals force adaptation

Mechanical forces, including pulling and pushing, are essential to deform the cell nucleus in order to facilitate movement into confinement^15,30,31,33^. Here, we examine the changes in nuclear shape relative to their spherical shape as a tool to infer whether the nucleus is “pulled” or “pushed” ^15,30–33^. The nuclear dimensions in the two unconfined directions are indicative of the forces applied onto the nucleus^41^. Using two-dimensional time-lapse microscopy data, we infer the 3D shape of the nucleus using the 2D aspect ratio and the measured nucleus Poisson ratio (Fig. 4a inset). This allows us to calculate the x-z aspect ratios (AR) of nuclei in the channel to those of force free nuclei (SI section 2). We introduce a dimensionless shape parameter *ε* = *AR_confined_*/*AR_free_*, where *AR_confined_* and *AR_free_* denote the confined and force free nuclear x-z aspect ratios, respectively. Values exceeding 1 indicate that in confinement the nucleus is being stretched in the direction of migration, while values below 1 indicate that the nucleus is compressed in the migration direction. As shown in Fig. 4d, in the CEM-regime *ε* initially rises with increasing confinement width up to a maximal value of 1.4 at a channel width of 12 µm. When transitioning into the CRM-regime, *ε* starts to decrease and eventually drops to values below 1 for channel widths below 7 µm, where it reaches a value of 0.5 at 4 µm confinement width. Thus, wild-type MDA-MB-231 cells show a non-monotonic dependence of the shape parameters *ε* with channel width, suggesting a change in the forces acting on the nucleus with varying confinement.

Based on our characterization of the mechanical properties of the nucleus in confinement, we can relate the differences in nuclear deformations to changes in the forces that are applied to the nucleus. We assume that there are two contributions to these forces: a pulling force *F_pull_* that is generated in front of the nucleus and a pushing force *F_push_* acting from behind the nucleus. Overall, these forces result in a total deformation force *F_deform_* = (*F_pull_* − *F_push_*)/2 on the nucleus in the direction of migration (SI section 2). If pulling forces dominate, there is a positive deformation force in the migration direction leading to an elongated nucleus (*ε* > 1), while a dominant pushing contribution results in a negative value of the deformation force with a compressed nucleus (*ε* < 1) (see Fig. 4d, inset). Thus, our mechanical nucleus model suggests that the change in the nuclear deformation behavior with changing channel widths can be interpreted as a transition from a pulling-dominated migration regime at wider channel widths to a pushing-dominated migration regime at channel widths below 7 µm.

### Mechanistic model for adaptive force generation

We explore the idea of an adaptive force generation mechanism by developing a model for cellular force generation during confined migration. The key aspect of the model is the combination of two confinement-dependent force generation mechanisms: pulling forces, generated by actomyosin contractility at the front of the cell, and pushing forces, regulated by the cortical tension in the rear (Fig. 5a). We incorporate these two mechanisms by generalizing a simple dynamical model that describes mesenchymal cell migration on 2D substrates ^35^ to 3D (see SI section 4 for details) and including nuclear deformations in confinement. Our model consists of three degrees of freedom: the nuclear position, the position of the leading protrusion of the cell, and a polarization (Fig. 5b). The nucleus and the protrusion are coupled through an elastic spring *k*. We absorb the friction coefficients of the nucleus and the protrusion into the spring constants and denote the rescaled spring constants as *k_n_* and *k_p_*, respectively. In previous work, we found that confinement of the protrusion can stimulate protrusion growth^35,42^, which we model by including a positive self-regulation on polarization in strong confinement (Fig. 5b(i)). Motivated by recent findings indicating that nuclear deformations can trigger increases in cortical tension through Ca^2+^ release^19,20^ (Fig. 5a(iii)), we thus include an increase in pushing forces with nuclear deformation (Fig. 5b(ii)). Further, we account for the confinement of the nucleus by increasing the effective nuclear drag coefficient *γ_n_*with decreasing channel width (Fig. 5b(iii)). Finally, we model the effect of elastic nuclear deformations by introducing an increasing elastic energy barrier with increasing nuclear confinement (Fig. 5b(iv)). To allow a comparison to our experimental analysis of the nuclear shapes, we use our mechanical model for nuclear deformations to relate the pulling forces, exerted by the protrusion onto the nucleus, and the pushing forces in our model to nuclear deformations.

**Figure 5:**
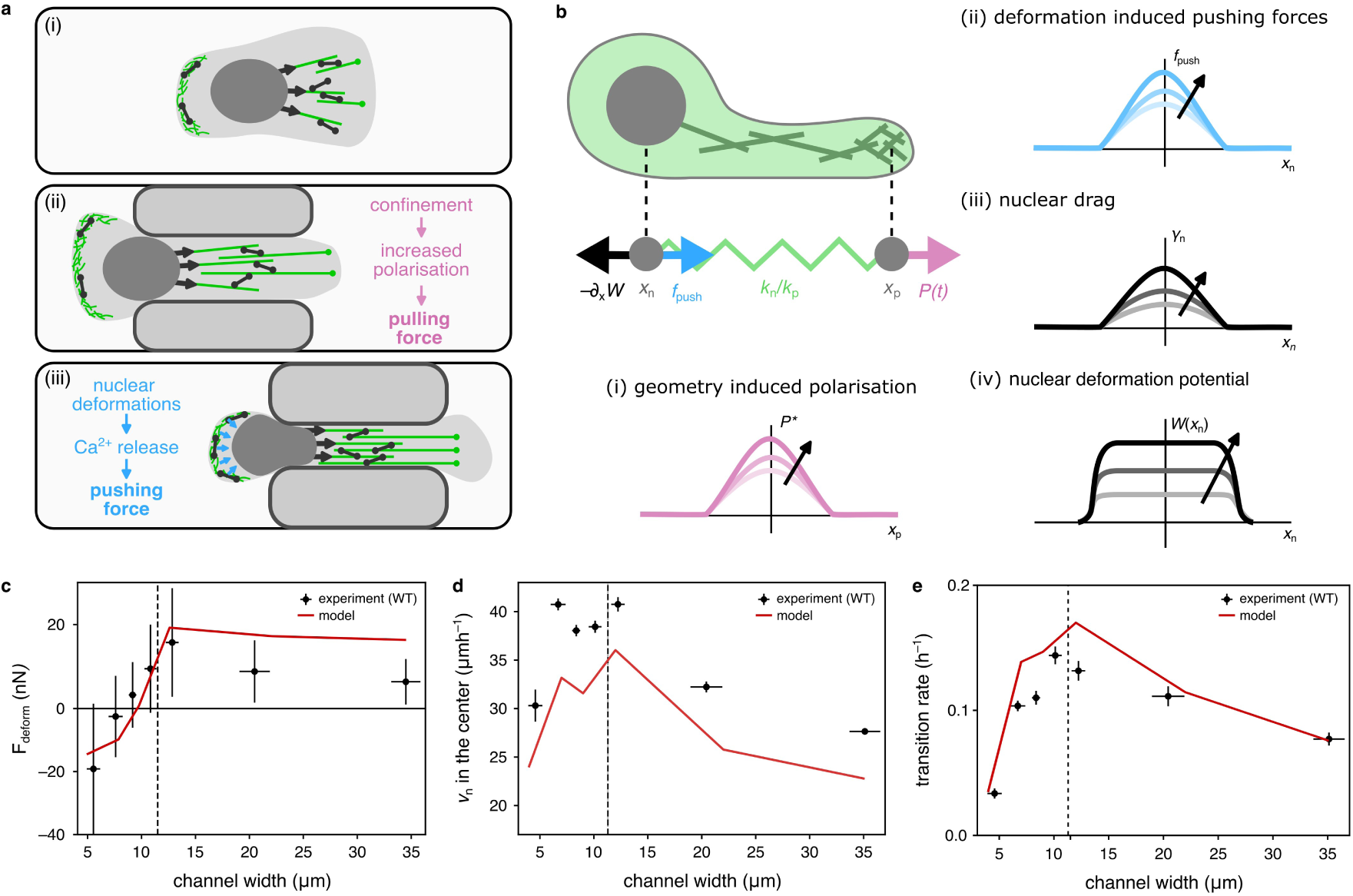
Mechanistic model for the adaptation of cellular forces to the degree of confinement. **(a)** Sketch of the proposed effect of the degree confinement on the cellular force production mechanisms. Compared to the unconfined case (i), confinement of the protrusion (ii) stimulates protrusion growth resulting in an increased pulling force. On the other hand, narrower protrusions contain less actomyosin, resulting in a reduction of the pulling force. Strong deformations of the nucleus (iii) trigger Ca^2+^ release, which results in a higher cortical contractility. Consequently, the pressure in the rear of the cell increases, resulting in an increased pushing force onto the nucleus. **(b)** Sketch of a minimal mechanistic model for mesenchymal cell migration. The protrusion and the nucleus are coupled elastically. Narrower protrusions result in a weaker elastic coupling between the nucleus and the protrusion (i). Protrusion growth is driven by a polarization force that accounts for the internal organization of the cell and that increases in confinement (ii). Additionally, when the nucleus gets deformed the cell can generate a pushing force that acts directly on the nucleus (iii) and the confinement induced deformations lead to a force that pushes the nucleus out of the channel (iv). **(c)-(e)** Fit of the mechanistic model to key experimental observations. Error bars: (c) Error bars represent one standard deviation. Error bars in (d), (e) associated with the y-axis represent the standard error, error bars of the channel width represent the standard deviation.

To constrain the free parameters of our mechanistic model, we simultaneously fit our model to a number of key experimental statistics: the deformation force acting in the direction of migration (Fig. 5c), the nuclear velocity in the center of the pattern (Fig. 5d) and the transition rate (Fig. 5e). Our model suggests the following interpretation of biphasic channel width dependence observed in both the nuclear deformations as well as the migration dynamics: At widths wider than the size of the nucleus (the CEM-regime), stronger confinement leads to enhanced polarization and thus protrusion growth. This results in a larger force pulling on the nucleus from the front resulting in increased nuclear velocities and transition rates (Fig. 5c- e). By contrast, for channel widths below the nuclear width (the CRM-regime), increased pushing forces from the rear of the cell result in a decreasing *F_pull_* − *F_push_*, eventually even reaching negative values (Fig. 5c). Despite the increased polarization and the additional pushing force, the increased nuclear friction result in a decrease of the predicted nuclear velocity in the channel (Fig. 5d). Additionally, the increasing elastic energy barrier, associated to nucleus deformations, hinders the movement of the nucleus into the channel, resulting in a drop of the transition rates (Fig. 5e).

To test the predictive power of our model after we have constrained all parameters, we calculate the probability distributions of the nuclear position and velocities in our simulations at different channel widths. The probability distribution of the nuclear position transitions from a broad distribution at wide channel widths, to a strongly double-peaked distribution for narrow channels, which agrees semi-quantitatively with our experiments (Fig. 2b, inset). Both in the experiment and the simulations, the distribution of nuclear velocities peaks at zero, irrespective of the channel width but decreases in spread with decreasing channel width (Fig. S13a, b). Further, our mechanistic model also allows us to connect features in the NCMs to cellular mechanisms. To do so, we compute the effective underdamped nuclear dynamics of our model (SI section 4). Similar to the dynamics inferred from experiments, we observe a deterministic flow from one chamber to the other with a pronounced acceleration in the channel (Fig. 3a, inset). Further, we find that our model successfully predicts most of the key features observed in the NCMs (Fig. 3b, inset and S13c, d): At the channel entrance the elastic barrier associated with nuclear deformations results in a region of Δ*F_w_* < 0, followed by a pronounced region of Δ*F_w_* > 0 associated with the recoil of the contractile actomyosin structures in the protrusion and the additional pushing forces acting onto the nucleus. Finally, as the cell leaves the channel, we observe a region of deceleration (Δ*F_w_* < 0) as the nucleus catches up with the protrusion. In conclusion, our mechanistic model demonstrates that cells transition from pulling to pushing dominated migration to generate sufficient deformation forces in confinement. This model not only explains the observed nuclear deformations but also allows for a mechanistic interpretation of the effective cellular dynamics inferred from experimental data.

### Perturbation of force generation using cytoskeleton inhibitors

A key conclusion of our analysis of the nuclear deformations is that cells can adaptively use two distinct force generation mechanisms during confined migration. To experimentally validate this, we aimed to disrupt the pulling and pushing mechanisms of the cells and examine the resulting impact on nuclear shapes. While both pulling and pushing forces are actin-myosin dependent, their mechanisms of force transmission are different. In particular, the generation of pulling forces requires the anchoring of stress fibers to the substrate, consequently leading to the formation of focal adhesions. To disrupt this process, we performed experiments in the absence of fibronectin coating. We observed a decrease in the relative contribution of pulling forces to the nuclear deformations (Fig. 4d, green line) (see Video S6). Both pushing and pulling forces rely on actomyosin contractility. To investigate how the balance between pulling and pushing forces is influenced by actomyosin contractility, we treat the cells with the ROCK inhibitor Y27632 (Fig. 4d, orange line) and the contractility activator Calyculin A (Fig. 4d, pink line) (see Videos S7 & S8). Our analysis suggests that in both Y27632 and Calyculin A treated cells, the relationship between pulling and pushing forces shifts in favor of pushing compared to the wildtype. Overall, we find that selective disruption of the pulling mechanism by targeting focal adhesions leads to changes in the observed nuclear deformation that are consistent with our mechanistic model (Fig. 4d).

## DISCUSSION

In summary, we studied the repeated self-imposed migration of single cells through compliant 3D channels. We found that in the regime of wide channels, with channel width wider than the nucleus (confinement-enhanced migration) transitions frequencies and velocities increases with confinement. In contrast, migration at sub-nucleus confinement (confinement-reduced migration) is impeded by 3D confinement. A similar biphasic behavior was observed in the elastic deformation of the nucleus from oblate to prolate shapes. Overall, we find that nuclear deformations are reversible with relative volume reductions of the nucleus up to 10%. The deformation has a multifaceted impact on the cell dynamics in the channel, with marked slowing down during the entry phase as reflected in the acceleration maps obtained through the inference of non-linear migration dynamics at varying channel widths. To explain these observations, we propose an extended dynamical cell migration model that accounts of the adaptive modulation of forces in response to confinement. The increase in pushing forces within confinement, together with elastic deformations of the nucleus and increased effective friction in the channel, explains both the observed nuclear deformations, as well as the overall migration dynamics of the cells across a broad range of channel widths.

Our dynamical model successfully predicts key features of the nuclear confinement maps, which allows for a mechanistic interpretation of the inferred dynamics in terms of confinement induced elastic nuclear deformations and cellular force adaptation. In the future, this approach could be further extended by using alternative inference approaches that account for additional degrees of freedom, allowing for the identification of other confinement adaptation mechanisms directly from data^43^.

While volume reduction of the nucleus due to the water-permeable nuclear membrane^29^ has been hypothesized, an experimental confirmation of the compressibility so far was difficult^28^. The observed decrease in nuclear volume within confinement, we report here, supports the notion of a purely elastic response with a distinct Poisson ratio. Previous work showed that some cell lines actively adapt their nuclear stiffness in confinement^44,45^. However, by comparing the Young’s moduli of confinement-deformed and undeformed nuclei probed by AFM, we found no evidence for nuclear mechanical adaptation effects, such as nuclear softening, at the time scale of our experimental system.

Both pulling and pushing forces have been qualitatively identified to play a role in confined cell migration^15,30–32^. Our analysis of nuclear deformations in confinement provides a quantitative measure of the balance between pulling and pushing forces, which indicates a confinement-induced adaptation of cellular force generation. A potential mechanism that could be involved in this force adaptation is the upregulation of cortical contractility in response to externally induced nuclear confinement^19,20^. We found that the pulling-dominated regime is reduced in the absence of focal adhesions, which agrees with a prior study wherein the disruption of focal adhesions at the cellular leading edge was achieved through laser ablation experiments^30^. By selective manipulation of the actomyosin activity, we showed that both up- and downregulation of actomyosin contractility led to a shift towards pushing dominated migration. A recent study classified cell transits of MDA-MB-231 cells through constrictions based on the cellular morphology^32^. Treatment with the contractility activator LPA resulted in a stronger rounding of the cell rear, while the contractility inhibitor Y27632 led to the opposite effect. This could indicate that the shift towards the pushing-dominated regime caused by the Y27632 treatment (Fig. 4D) might not be driven by an increase in pushing forces but an even stronger decrease in pulling forces. Further research is required to experimentally verify the potential mechanism of increased cortical contractility through intracellular calcium imaging and identify whether this mechanism is universally applicable or specific to certain highly migratory cell lines, such as cancer cells.

Overall, our study shows that during self-imposed confined cell migration, the nucleus can be considered as a compressible elastic object with a Poisson ratio of 0.40 ± 0.02, independent of the channel width, which is translocated by a responsive cytoskeleton that is sensitive to confinement. Previous studies^19,20^ have shown that cells use their nucleus to sense and adapt to externally imposed confinement. Our work adds to this by providing evidence that the force generating cytoskeleton adapts by switching from pulling to pushing forces to overcome constrictions. This contributes to a more comprehensive mechanistic understanding of the complex interplay between confinement, the nucleus, and the cytoskeleton during mesenchymal cell migration.

## Methods

### Cell culture

MDA-MB-231 human breast carcinoma epithelial cells, co-expressing histone-2B mCherry (gift from Timo Betz, Göttingen), were cultured in a standard common growth medium, specifically L-15 (Sigma), supplemented with 10% fetal bovine serum (Sigma). The cells are cultivated at a temperature of 37°C until reaching a confluence level of 80-90%. Following this, the cells were washed and trypsinized for 4min. For experimental purposes, the cell solution was centrifuged at 1,000 r.c.f. for 3min. Following this, the cells were re-suspended in L-15 medium. Approximately 20,000 cells were seeded per µ-dish (ibidi) and allowed to adhere for a minimum duration of 3h. For inhibitor experiments, 0.5 nM Calyculin A (Thermo Fisher), 30 μM Y27632 (Sigma Aldrich) was added 2 hours before the start of the experiment. For live cell imaging of actin, MDA-MB-231 cells stably transduced with LifeAct GFP and histone-2B mCherry were used (gift from Timo Betz, Göttingen).

### Preparation of dumbbell-shaped hydrogel cavities

The experimental procedure involved the use of Polydimethylsiloxan (PDMS) stamps mixed in a 10:1 monomer cross-linker ratio, poured on the specific silicon wafer, degassed and cured over night at 50°C. PDMS stamps containing small, squared pillars (200x200x20 µm³, see Fig. S1) were activated for 3min in a UV cleaner (PSD-UV, novascan). Following activation, the stamps were stamped on a petri dish (µ-Dish ibiTreat, ibidi). A hydrogel precursor droplet was put next to the stamps and gets drawn into the free space between the stamps provided by the small pillars. The hydrogel precursor was then photopatterned using the PRIMO module (Alvéole) integrated into an automated inverted microscope (Nikon Eclipse Ti). The placement of the desired dumbbell pattern was achieved using the Leonardo software (Alvéole), and subsequently, it was exposed to UV-light with a dose of 3 mJ/mm² to induce polymerization of the hydrogel. After the illumination process, the stamps were removed, and the dish thoroughly washed with milliQ water and rehydrated with PBS for 5min. Following rehydration, the dish was incubated with 50 μg ml^-1^ human fibronectin (YO Proteins) solution for 45 min and subsequently rinsed with PBS. After waiting for 45 min, the dish was subsequently rinsed with PBS and stored in PBS at room temperature until cells were seeded on it.

### Microscopy and cell tracking

Measurements were performed in time-lapse mode for up to 40h on a Nikon-Eclipse Ti-E inverted microscope. To ensure consistent incubation conditions throughout the measurements, the microscope is equipped with gas incubation and a heating system (Okolab). Brightfield and fluorescence images of the fibronectin-coated pattern and the co-expressed labeled histones were acquired every 10min. In order to enhance the quality of the nuclei images, a bandpass filter was applied. Following this, the images were binarized, and the positions of the nuclei’s center-of-mass were determined using the Analyze Particles plugin in ImageJ.

### Nuclear volume analysis

To analyze the changes in nuclear volume over time, z-stacks within the frame of a timelapse recording over 20h z-stacks of MDA-MB-231 cells transmigrating through the constriction were captured with a confocal microscope (LSM-980, Zeiss). To provide standard incubation conditions throughout the measurements, the microscope is equipped with gas incubation and a heating system (Okolab). The nuclear volume for each timestamp was analyzed by the software ‘arivis4D’ (Zeiss) (see Fig. S2). For further details, see Supplementary Section 1.

### Stiffness evaluation via AFM

Atomic force microscopy was employed to assess mechanical properties, including the Young’s modulus, of both the hydrogel and the cell nuclei. The measurements were conducted using a JPK NanoWizard II (JPK Instruments), which was interfaced with an inverted optical microscope (Zeiss, Axiovert 200M). The utilized cantilevers were modified with beads, possessing diameters of either 3.5 *µm* (NanoAndMore GmbH; type: CP-PNPL-SiO-B, nominal spring constants of 0.08 *N/m*) or 3.6 *µm* (sQUBE, type: CP-CONT-PS-B, nominal force constants between 0.02 and 0.77 *N/m*). To calibrate the cantilever sensitivity, the slope of the force-distance curve against the hard petri dish substrate was recorded. Subsequently, the spring constants of the cantilevers were measured using the thermal noise method provided by the AFM software (JPK SPM). The Young’s modulus of the hydrogel was determined by applying large forces (>10 nN) through the bead, resulting in indentation depths of up to 2 *µm*. Likewise, the stiffness of the cell nuclei was determined using the same atomic force spectroscopy method by applying force (5 nN) to the cell nuclei (see Fig. S3b). Typically, a square grid consisting of 64 points with 1 *µm* spacing between neighbors was tested at a single position to ensure statistically robust results. Force-distance curves were obtained with an extension speed of 2.5 *µm/s.* The Young’s moduli of the tested objects were extracted by fitting the force-distance curves to a hertz contact model.

### Bead displacement analysis

The tracking of fluorescent nano-beads embedded within the hydrogel and the cell nuclei was carried out using the TrackPy Python package. In order to observe the movement of a particle, its displacement relative to a neutral reference position was calculated. For quantitative analysis of the observable deformation in the gel, the tracking algorithm was employed to trace the positions of the fluorescent nano-beads. To visualize their displacement, the difference between the positions of the marker beads at each time step and their initial neutral position was determined. Considering that the marker beads are embedded within the gel, this is due to their larger diameter of 200 nm compared to the approximate mesh size of the hydrogel, which is approximately 40 nm. Consequently, the displacement field exhibited by the beads reflects the deformation field of the surrounding hydrogel. For further details, see Supplementary Section 1.

### Nuclear shape analysis

To quantify the effect that external forces have on the confined nucleus in the center of the channel, we compute the deviation from an isotropic (force free) expansion of the nucleus under compression through the channel walls. To find an approximate expression for the dimensions of the nucleus under compression in the presence of pulling and pushing forces acting along the *x*–direction, we first calculate the shape of the unconfined nucleus with pulling and pushing forces applied and then apply the confinement induced deformation in the *y*-direction together with an isotropic expansion in the *x* /*z*-direction. For further details, see Supplementary Section 2.

## Acknowledgements

We thank D. Brückner, T. Brandstätter, B. Sabass, J. Guck, P.Ronceray and C. Cykes for helpful discussion and C. Leu for the preparation of wafers. This work was funded by the Deutsche Forschungsgemeinschaft (DFG, German Research Foundation) – Project-ID 201269156 – SFB 1032 Projects B01 and B12).

## Author contributions

S.S. and J.O.R. designed experiments; S.S., M.M.K. and M.B. performed experiments; J.F., S.S. and M.M.K. analyzed data; J.F. and C.P.B. developed the theoretical model and performed the inference. S.S., J.F., C.P.B. and J.O.R. wrote the manuscript.

## Reporting summary

Further information on research design is available in the Nature Portfolio Reporting Summary linked to this article.

## Data availability

Experimental and simulation data is available from the corresponding author upon request.

## Code availability

Python code to analyze data and perform numerical simulations is available from the corresponding author upon request.

