## Supplementary Information for "Nuclear deformation and dynamics of migrating cells in 3D confinement reveal adaptation of pulling and pushing forces"

#### Contents

**0. Movie descriptions** ........ **- 3 -**

**1. Further experimental details** ........ **- 4 -**

1.1. Hydrogel precursor ........ - 4 -

1.2. Photolithography ........ - 4 -

1.3. Microstructure design ........ - 7 -

1.4. Cell exclusion criteria ........ - 7 -

1.5. Nuclear growth analysis ........ - 7 -

1.6. Determination of the Poisson ratio ........ - 8 -

1.7. Stiffness evaluation via AFM ........ - 9 -

1.8. Measured normal forces exerted by the cell nucleus ........ - 10 -

1.9. Analysis of bead displacement orientation ........ - 11 -

**2. Nuclear shape analysis** ........ **- 12 -**

2.1. Theoretical model for the normal forces ........ - 12 -

2.2. Calculation of the shape parameter ........ - 13 -

2.3. Deducing the force balance on the nucleus ........ - 15 -

2.4. Plastic deformations and shape recovery ........ - 17 -

**3. Inference of the dynamical signature of the confinement** ........ **- 18 -**

3.1. Symmetries of the basis functions ........ - 18 -

3.2. Criteria for the model selection ........ - 19 -

3.3. Uniqueness of the inferred models ........ - 20 -

**4. Mechanistic model** ........ **- 23 -**

4.1. General structure of the model ........ - 23 -

4.2. Nuclear friction ........ - 24 -

4.3. Deformation barrier ........ - 25 -

4.4. Pushing force ........ - 25 -

4.5. Implementation and simulations ........ - 27 -

4.6. Calculation of the effective underdamped dynamics ........ - 27 -

4.7. Comparison of experimental data and model predictions ........ - 29 -

**References** ........ **- 31 -**

### 0. Movie descriptions

**Supplementary Video 1**
Single MDA-MB-231 cell transitioning repeatedly between the square islands of the dumbbell-shaped cavities. Transitions are usually preceded by the formation of a protrusion along the bridge. The cells are co-expressing histone-2B mCherry and F-actin (LifeAct-TagGFP2) to allow the visualization of the nucleus and actin distribution. The bridge width shown in movie 1 is $W$ = 16 µm. Scale bar: 20 µm.

**Supplementary Videos 2-3**
Single MDA-MB-231 cells transitioning repeatedly between the square islands of the dumbbell-shaped cavities. Transitions are usually preceded by the formation of a protrusion along the bridge. The cells are co-expressing histone-2B mCherry to allow automated tracking of cell positions. The bridge widths shown in movies 2 and 3 are $W$ = 18 and 5 µm respectively. Scale bar: 20 µm.

**Supplementary Video 4**
4D experiment of a single MDA-MB-231 cell transitioning through the thin bridge. Cell nucleus is shown in blue and embedded fluorescent-nanobeads are shown in red. The cell is co-expressing histone-2B mCherry, enabling the determination of the nuclear volume at each time point.

**Supplementary Video 5**
Single MDA-MB-231 cell transitioning through the thin bridge. Cell nucleus and embedded fluorescent-nanobeads are shown in grey. The hydrogel composition was chosen soft to allow recognizable bead displacement due to the nucleus squeezing through the constriction. The length and colour of the arrows represent the amount of displacement from their neutral position and the direction in which the arrow shows indicates the direction in which the deformation went from its neutral position. Scale bar: 20 µm.

**Supplementary Video 6**
Single MDA-MB-231 cell transitioning repeatedly between the square islands of the dumbbell-shaped cavities without fibronectin coating. Transitions are usually preceded by the formation of a protrusion along the bridge. The cell is co-expressing histone-2B mCherry to allow automated tracking of the cell position. The bridge width shown in movie 6 is $W$ = 8 µm. Scale bar: 20 µm.

**Supplementary Video 7**
Single MDA-MB-231 cell treated with the ROCK-inhibitor Y27632 transitioning repeatedly between the square islands of the dumbbell-shaped cavities. Transitions are usually preceded by the formation of a protrusion along the bridge. The cell is co-expressing histone-2B mCherry to allow automated tracking of the cell position. The bridge width shown in movie 7 is $W$ = 15 µm. Scale bar: 20 µm.

**Supplementary Video 8**
Single MDA-MB-231 cell treated with the myosin activator Calyculin A transitioning repeatedly between the square islands of the dumbbell-shaped cavities. Transitions are usually preceded by the formation of a protrusion along the bridge. The cell is co-expressing histone-

2B mCherry to allow automated tracking of the cell position. The bridge width shown in movie 8 is $W = 9$ µm. Scale bar: 20 µm.

**Supplementary Video 9**
Exemplary z-stack of the microstructures made of PEG-NB hydrogel captured by a confocal microscope. GFP-labelled Dextran (10 kDa) is used to analyze the structures as function of the height $z$ as the Dextran is not able to diffuse into the hydrogel due to the molecular size. Scale bar: 40 µm.

### 1. Further experimental details

#### 1.1. Hydrogel precursor

The precursor solution for hydrogel formation was prepared using phosphate-buffered saline (PBS) containing 5 mM of 20 kDa 8-armed PEG-norbornene (PEG-NB, JenKem Technology). To enable crosslinking, an off-stoichiometric amount (0.4 – 0.8) of 1 kDa PEG-dithiol (Sigma-Aldrich), was added along with 3 mM of the photo-initiator lithium phenyl-2,4,6-trimethylbenzoylphosphinate (LAP, Sigma-Aldrich). The inclusion of the PEG-dithiol cross-linker and LAP photo-initiator enables the formation of a stable crosslinked network within the hydrogel. The mechanical properties of the resulting hydrogel can be directly influenced by varying the quantity of Peg-NB monomers and/or the amount of cross-linker. The cross-link ratio $r_c$ is defined as the ratio of functional groups present in the cross-linker (two thiol groups per cross-linker) to the functional groups present in the PEG-NB monomer (eight norbornene groups per monomer).

$$r_c = \frac{2c(\text{dithiol})}{8c(\text{PEG-NB})} \tag{S1}$$

Fluorescent nano-beads, characterized by a size of 0.2 µm, were incorporated into the precursor solution with a total concentration of 5.8E+08 $ml^{-1}$ (Distrilab). These embedded nano-beads function as force sensors due to the elastic properties of the hydrogel.

#### 1.2. Photolithography

Silicon wafers undergo a series of treatment steps to prepare them for further processing. Initially, the wafers are subjected to hydrofluoric acid treatment (TECNIC) to clean the surface. Following this, a layer of photoresist AZ40XT (AZ Electronic Materials) is spin coated onto the wafer, resulting in a thickness of approximately 20 µm. The coated wafer is then subjected to a soft bake process. To achieve the desired patterns, laser direct imaging (Protolaser, LPKF) is used. Following exposure, a post-exposure bake step is performed to further stabilize the resist. Subsequently, the wafer undergoes development using AZ 726 MIF (AZ Electronic Materials) to remove the unexposed photoresist, revealing the patterned geometries. In the final step, the wafer surface is treated with (Trichloro(1H,1H,2H,2H-perfluoro-octyl)silane (Sigma-Aldrich) in a process known as silanization. This treatment imparts a hydrophobic character to the wafer surface by bonding the silane molecules, which helps control surface interactions in subsequent processes.

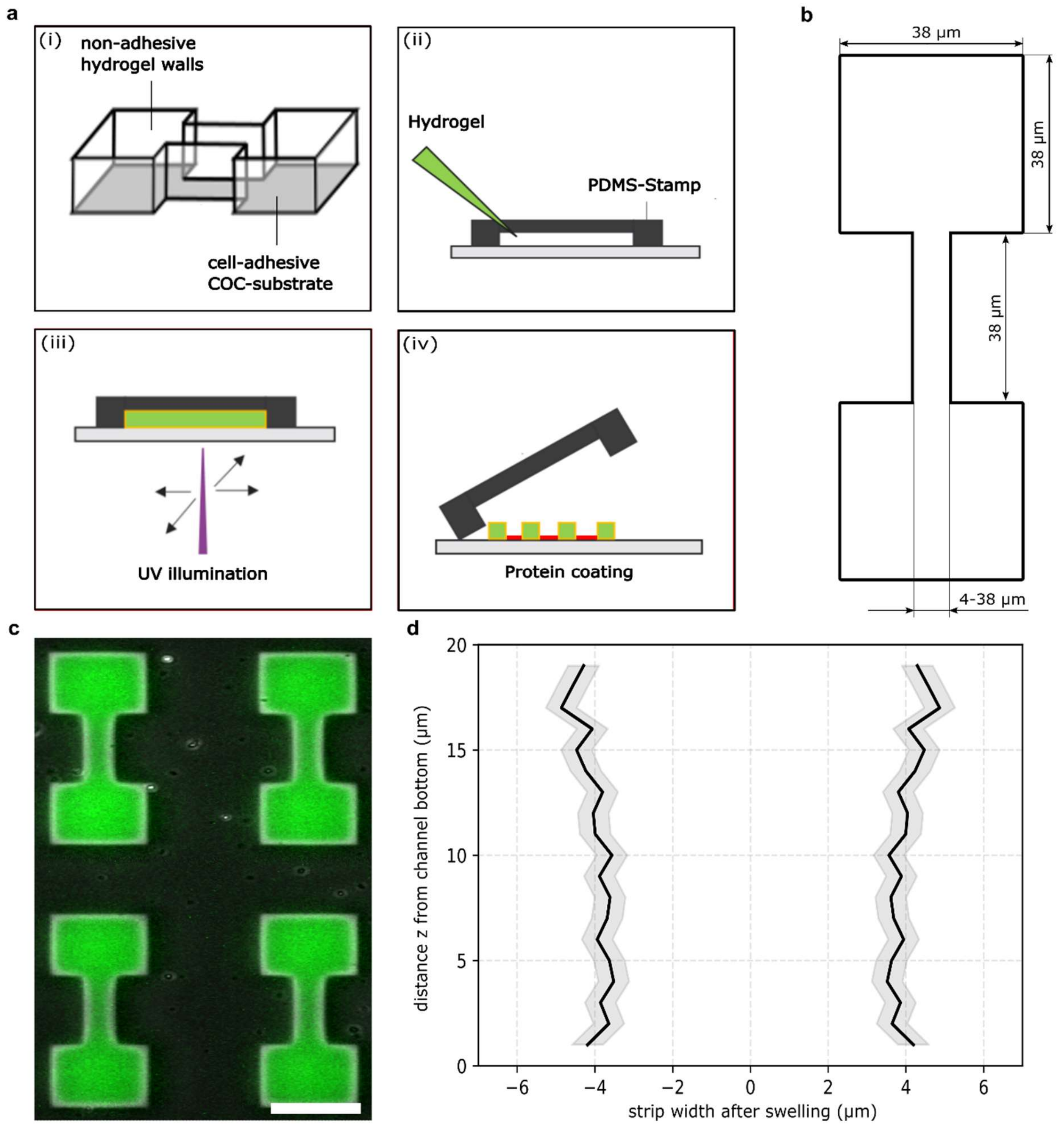


**Figure S1: Validation of the experimental specifics. (a)** Step-by-step protocol for fabricating hydrogel-dumbbell cavities. (**b)** Determination of the characteristic dimensions of the dumbbell structures, with the bridge width serving as the variable parameter. **(c)** Illustration of the fibronectin-coated bottom surface of the dumbbell structures (scale bar, 40 um). **(d)** Analysis of the correlation between the constriction width and the spatial distance *z* from the bottom of the channel (see Video S9).

| Cell line | Bridge width (µm) | Cells | Transitions | Timepoints |
|---|---|---|---|---|
| MDA-MB-231 wildtype | 4.6 ± 0.65 | 144 | 82 | 16468 |
| | 6.7 ± 0.7 | 333 | 639 | 41127 |
| | 8.4 ± 0.4 | 211 | 434 | 26256 |
| | 10.1 ± 0.6 | 164 | 449 | 20770 |
| | 12.2 ± 0.6 | 102 | 279 | 13939 |
| | 20.4 ± 1.5 | 99 | 206 | 12351 |
| | 35.1 ± 1.4 | 249 | 318 | 27760 |
| MDA-MB-231 Y27632 | 4.3 ± 0.9 | 125 | 9 | 18101 |
| | 6.6 ± 0.7 | 237 | 85 | 30516 |
| | 8.5 ± 0.4 | 142 | 73 | 16333 |
| | 10.3 ± 0.7 | 115 | 98 | 14197 |
| | 12.1 ± 0.5 | 109 | 104 | 14144 |
| | 16.0 ± 0.9 | 153 | 133 | 20006 |
| | 19.7 ± 0.9 | 49 | 55 | 7005 |
| MDA-MB-231 w/o FN | 7.1 ± 0.6 | 24 | 28 | 3725 |
| | 8.6 ± 0.3 | 43 | 37 | 6324 |
| | 10.0 ± 0.6 | 46 | 61 | 5695 |
| | 12.8 ± 0.8 | 52 | 38 | 6999 |
| | 15.0 ± 0.8 | 117 | 131 | 15957 |
| | 19.4 ± 0.8 | 45 | 72 | 6096 |
| MDA-MB-231 Calyculin A | 6.8 ± 0.8 | 41 | 67 | 6007 |
| | 8.4 ± 0.3 | 29 | 57 | 3439 |
| | 10.2 ± 0.6 | 23 | 75 | 2672 |
| | 12.1 ± 0.7 | 12 | 37 | 1535 |
| | 19.2 ± 1.1 | 41 | 29 | 7005 |

**Table S1: Bridge widths *w* of the micro-cavities and number of statistics collected. Number of cell trajectories and number of transitions recorded for each bridge width *w*.**

#### 1.3. Microstructure design

All dumbbell-shaped cavities were specifically designed to feature two islands of equal size ($(37.5 \pm 0.9)^2\ \mu m^2$) connected by a bridge of uniform length ($(35.0 \pm 1.0)\ \mu m$). The width of the bridge denotes as *w*, and the corresponding number of cell trajectories for each value of *w* are presented in Table S1. The micropattern without constriction was designed to possess a similar width and total length as the two-state pattern with L = 35 µm, and thus has dimensions (($109.0 \pm 2.0$) µm) x ($37.0 \pm 1.5$) µm).

#### 1.4. Cell exclusion criteria

We track the position of many cells to determine the transition statistics of cells migrating through constrictions with varying bridge width. To limit the effects of ambiguous or abnormal migration behaviour, we apply the following inclusion criteria in the analysis of migration in our hydrogel micro-cavities.

1. Only a single cell occupies the micro pattern. Trajectories are cut when the cell rounds up for division.
2. The cell and its protrusions are entirely confined within the borders of the microstructure.
3. Throughout the entirety of the experiment, the cell shows no abnormalities such as the presence of multiple nuclei, cell death, or detachment from the substrate.
4. Cell dynamic statistics are included when a protrusion forms and extends to the second adhesion-site, even if the cell is unable to transmigrate through the constriction.

Criteria 1-3 represent fundamental conditions for single cell experiments within micro-cavities, and these criteria alone are applied to cells undergoing migration in cavities without constriction. Criterion 4, however, is exclusive to our dumbbell-shaped hydrogel cavities, as the migratory potential of cells within these structures is heavily influenced by the width of the constricted region, particularly in terms of physical confinement. Despite the inability of cells to transmigrate through these narrow constrictions, we consider these cells in our analysis, as such behaviour is deemed unremarkable in this context.

#### 1.5. Nuclear growth analysis

For nuclear volume analysis, data were imported into the Arivis Vision 4D software. Subsequently, each dataset underwent initial bleach correction. Following this correction, individual nuclei were segmented and size-filtered to eliminate extraneous noise components. The segmented objects were subjected to manual proofreading, and parameters were adjusted as required, enabling the calculation of nuclear volume within the software.

To investigate the overall trend of increasing nuclear volume over time (Fig. 4a), we performed control experiments of MDA-MB-231 cells caught within a big sized hydrogel cavity without constriction ($(50.0 \pm 2.0)^2\ \mu m^2$) (see Fig. S2a). The evaluation of the nuclear volumes reveals no significant differences in the nuclear growth rate between the confined and unconfined conditions (mean ± SD for n = 3 cell nuclei per condition). Thus, the increasing nuclear volume over time originates from the cell growth.

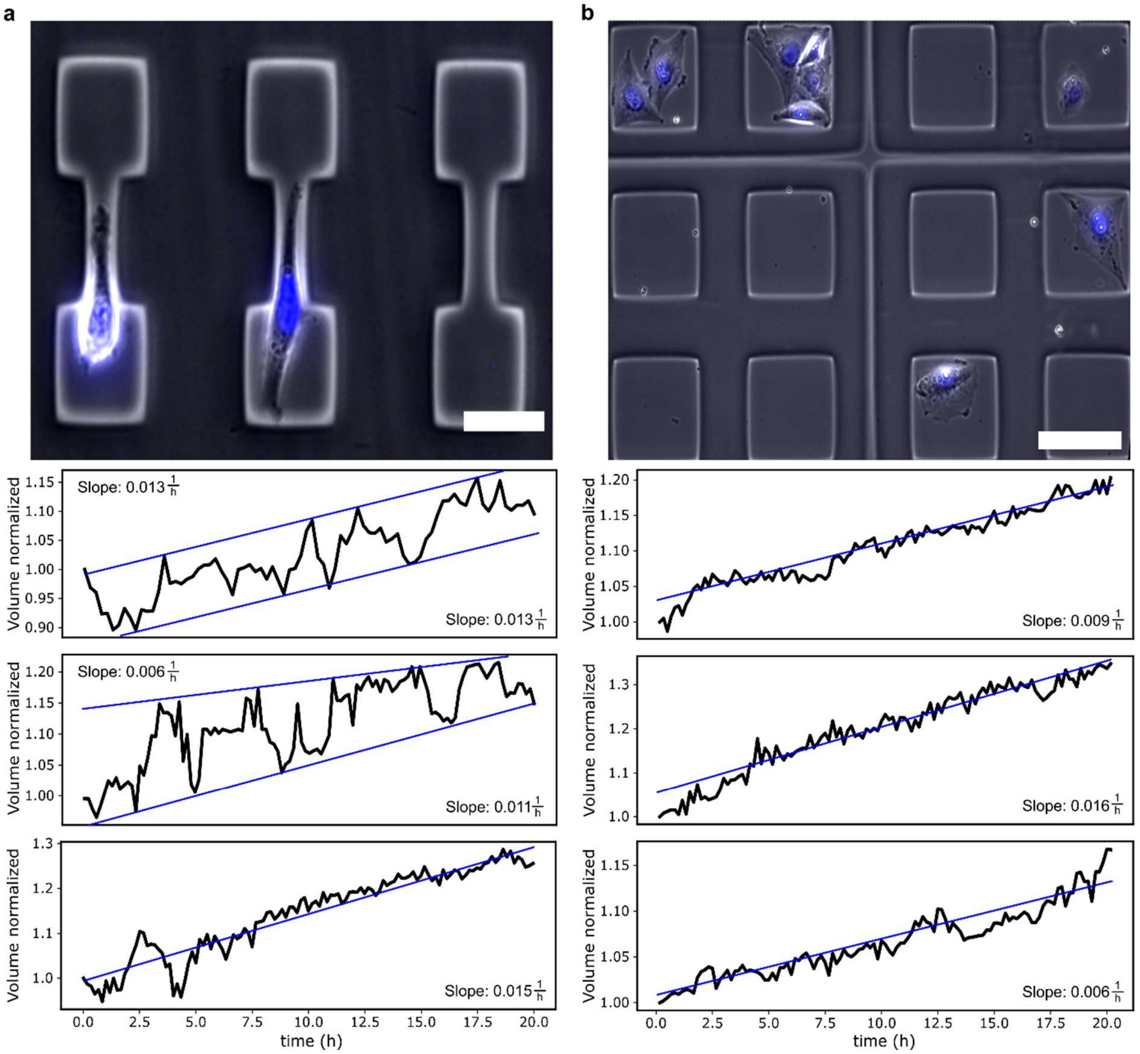


**Figure S2: Analysis of nuclear volume alterations over time. (a)** Cells located within dumbbell-shaped cavities. The normalized cell volume exhibits significant variations depending on the nuclear position within the dumbbell structure (scale bar, 40 µm). **(b)** Cell nuclei positioned within hydrogel cavities where cells do not encounter constrictions. The normalized cell volume exhibits progressive growth over time (scale bar, 50 µm).

#### 1.6. Determination of the Poisson ratio

The change in volume of a material under compression can be characterized in terms of the Poisson ratio $\nu$. A value of $\nu = 0.5$ characterizes an incompressible material, while values below 0.5 indicate that the material's volume changes under compression. To determine the Poisson ratio of the cell nucleus $\nu_n$, we consider the changes in volume and width of nuclei entering the constriction. The Poisson ratio $\nu_n$ is given by[1]

$$\nu_n = \left(\frac{1}{2} - \frac{\Delta V\, W}{2V\, \Delta \mathrm{W}}\right). \quad \text{(S2)}$$

With the following derivatives of $\nu$ with respect to $V_{rel} = \frac{\Delta V}{V}$ and $W_{rel} = \frac{\Delta \mathrm{W}}{W}$

$$\frac{d\nu_n}{dV_{rel}} = \left(-\frac{1}{2\,W_{rel}}\right) \tag{S3}$$

and

$$\frac{d\nu_n}{dW_{rel}} = \left(\frac{\mathrm{V_{rel}}}{2\,W_{rel}^2}\right), \tag{S4}$$

we then calculate the standard deviation (*SD*) of $\nu_n$, which is given by

$$SD_{\nu_n} = \sqrt{\left(\frac{\mathrm{V_{rel}}}{2\,W_{rel}^2}\,SD_{W_{rel}}\right)^2 + \left(-\frac{1}{2\,W_{rel}}\,SD_{V_{rel}}\right)^2}. \tag{S5}$$

#### 1.7. Stiffness evaluation via AFM

AFM experiments have been performed to measure the stiffness of the hydrogel and the cell nuclei. The median of the Young's modulus for the soft hydrogel was measured at 3 kPa over 1465 positions in more than 20 different samples and the hard hydrogel was measured at 24 kPa over 256 positions in four different samples.

To determine the nuclear stiffness of MDA-MB-231 cells, cells were adhered on a petri dish (µ-Dish ibiTreat, ibidi) without any confinement. The median of the Young's modulus for the cell nuclei was measured at 368 Pa. These measurements include 1920 different positions on 20 different cells. The Young's modulus of the MDA-MB-231 cells agrees well with literature values[2,3].

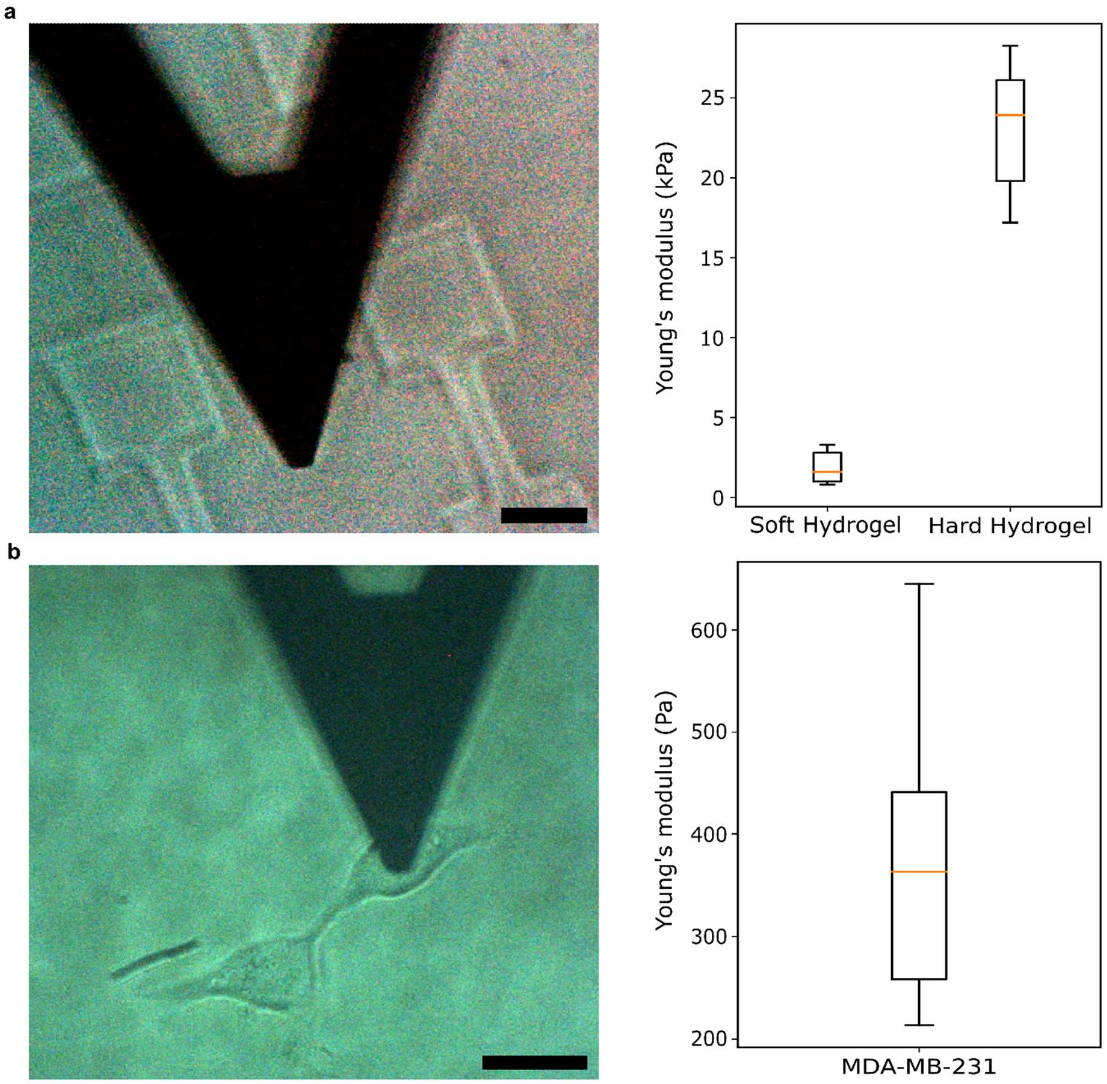


**Figure S3: Stiffness measurements of hydrogel and cell nuclei performed with an AFM. (a)** Evaluation of hydrogel stiffness by measuring the interaction between two dumbbell structures, considering various compositions of the hydrogel (Box plot ($n_{soft}$ = 1465; $n_{hard}$ = 256) with whiskers extending to ± 1.5 × IQR) (scale bar, 20 µm.) **(b)** Stiffness measurement of the nuclei in MDA-MB-231 cells (Box plot (n = 1920) with whiskers extending to ± 1.5 × IQR) (scale bar, 20 µm).

#### 1.8. Measured normal forces exerted by the cell nucleus

In order to evaluate the measured normal forces exerted by the nucleus, a classical Hertz' model for the gel deformation was assumed on the basis of the displacement of the fluorescent nano-beads that follow the deformation of the hydrogel. The hydrogel pushes back against an indenting sphere with the Hertz' force, according to the following expression[4,5]:

$$f_h(\delta) = \frac{4}{3}\frac{E_h}{1-{\nu_h}^2}\sqrt{R_i}\,\delta^{3/2} \qquad \text{(S6)}$$

Here, $\delta$ describes the indentation into the hydrogel wall, and $R_i$ the initial radius of the (unsqueezed) cell nucleus. $E_h$ and $\nu_h$ are the Young's modulus and Poisson's ratio of the hydrogel, respectively.

#### 1.9. Analysis of bead displacement orientation

Analyzing the hydrogel's motion when the cell nucleus is completely enclosed within the channel, with particular focus on the orientation of the arrows, provide valuable insights into the influence of friction within our system. The angle was determined by considering the y- and x-values of the displacement arrow.

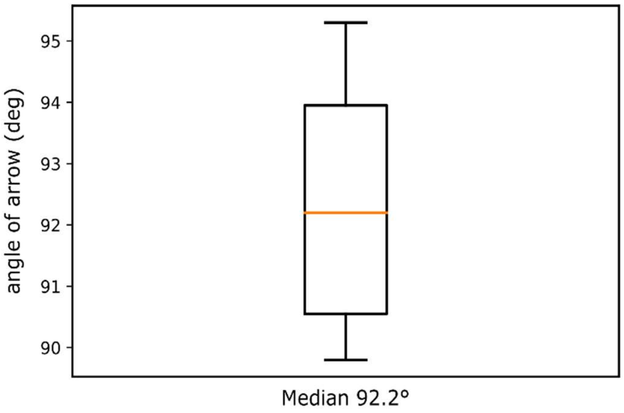


**Figure S4: Angle of bead displacement:** The median value of displacement along both the vertical and horizontal axes relative to the hydrogel wall surface (Box plot (n = 500) with whiskers extending to ± 1.5 × IQR).

### 2. Nuclear shape analysis

#### 2.1. Theoretical model for the normal forces

To calculate the expected indentation of the hydrogel by a passing cell, we use a simple Hertz model, as a soft sphere is compressed in between two soft half-spaces (see Fig. S5). As the sphere and constricting material are in equilibrium, one can assume that the hydrogel wall is exposed to a force similar to that of a hard sphere and that at the same time, the nucleus behaves like a sphere that is squeezed between similarly infinitely hard walls.

In conclusion, the force exerted by the nucleus squeezed by a length of $\alpha$ is given by the following equation[5]:

$$f_n(\alpha) = \frac{4}{3}\,\frac{E_n}{1-{\nu_n}^2}\,\sqrt{R}\,\alpha^{3/2} \tag{S7}$$

And the force that the hydrogel pushes back with against a sphere indenting $\delta$ into it, $F_h$ is given by Equation (S6). Here, $R_i$ describes the initial radius of the cell nucleus (unsqueezed), and R is the actual radius when squeezed. Equating Eq. (S6) and (S7) yields the expected equilibrium displacement (see Fig. S5d). For this, we write the indentation of the hydrogel as $\delta = \left(\frac{h}{2} - \frac{h_i}{2}\right)^{3/2}$ and the length of the squeezed nucleus as $\alpha = \left(R_i - \frac{h}{2}\right)^{3/2}$. Taken together, we get the following equation:

$$\frac{4}{3}\frac{E_n}{1-\nu_2^2}\sqrt{R_i}\left(R_i - \frac{h}{2}\right)^{3/2} = \frac{\left(\frac{E_n}{1-\nu_n^2}\right)^{\frac{2}{3}}}{\left(\frac{E_h}{1-\nu_h^2}\right)^{\frac{2}{3}} + \left(\frac{E_n}{1-\nu_n^2}\right)^{\frac{2}{3}}}\left(R_i - \frac{h_i}{2}\right). \tag{S8}$$

We consider the mechanical properties of the hydrogel and nucleus, respectively. The Young's modulus and Poisson's ratio of the hydrogel are denoted as $E_h$ and $\nu_h$, while the Young's modulus and Poisson's ratio of the nucleus are denoted as $E_n$ and $\nu_n$. The initial height of the constriction, represented as $h_i$, refers to the distance between the hydrogel walls when no cell is present. Similarly, the initial radius of the cell nucleus, denoted as $R_i$, represents its size when it is not subjected to a constriction.

When a cell is located within the constriction, the distance between the walls $h$ or the diameter of the nucleus changes. If the hydrogel and the nucleus have similar stiffness properties, and the constriction width is sufficiently small, a significant difference between the initial height $h_i$ and the final height $h$ can be observed. This difference occurs due to the displacement of the hydrogel caused by a displacement value $\delta$. The displacement can be calculated by

$$\delta = \frac{h}{2} - \frac{h_i}{2} = \frac{\left(\frac{E_n}{1-\nu_n^2}\right)^{\frac{2}{3}}}{\left(\frac{E_h}{1-\nu_h^2}\right)^{\frac{2}{3}} + \left(\frac{E_n}{1-\nu_n^2}\right)^{\frac{2}{3}}}\left(R_i - \frac{h_i}{2}\right). \tag{S9}$$

This equation describes the expected deformation of the hydrogel when a cell nucleus is squeezed inside a hydrogel constriction.

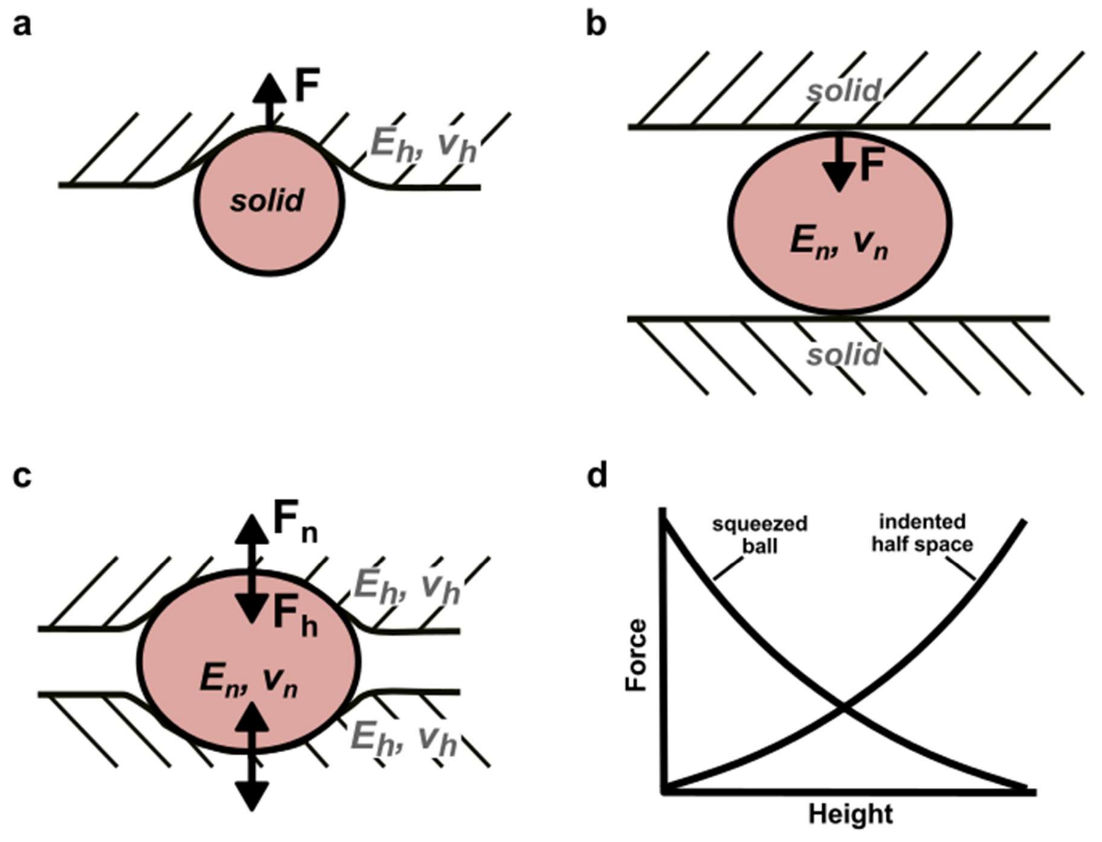


**Figure S5: Illustrations of the relevant contact mechanics setups. (a)** A solid sphere is pushed into an elastic half-space. **(b)** An elastic sphere experiences compression when confined between two rigid boundaries. **(c)** A deformable sphere undergoes squeezing between two elastic half-spaces. **(d)** An illustrative depiction of the anticipated state of equilibrium between a soft sphere and a flexible half-space.

#### 2.2. Calculation of the shape parameter

To quantify the effect that external forces have on the confined nucleus in the centre of the channel, we compute the deviation from an isotropic (force free) expansion of the nucleus under compression through the channel walls. To find an approximate expression for the dimensions of the nucleus under compression in the presence of pulling and pushing forces acting along the $x$-direction (see Fig. S6), we first calculate the shape of the unconfined nucleus with pulling and pushing forces applied and then apply the confinement-induced deformation in the $y$-direction together with an isotropic expansion in the $x$-/$z$-direction.

The sum of pushing and pulling forces induce a strain $u_{xx}$ in the $x$-direction. The strains in the orthogonal directions are then given by[6] $u_{yy} = -v_n u_{xx}$ and $u_{zz} = -v_n u_{xx}$. For small deformations along the $x$-direction, the strains can be written as[7] $u_{xx} = \frac{dx}{x}$, $u_{yy} = \frac{dy}{y}$, and $u_{zz} =$

$\frac{dz}{z}$. To find deformations $\Delta y$ and $\Delta z$ induced by the forces acting in $x$-direction, we integrate the infinitesimal strains and use that $u_{yy} = u_{zz} = -\nu_n u_{xx}$, such that

$$\int_{y_0}^{y_0+\Delta y} \frac{dy}{y} = \int_{z_0}^{z_0+\Delta z} \frac{dz}{z} = -\nu_n \int_{x_0}^{x_0+\Delta x} \frac{dx}{x}, \tag{S10}$$

from which we get

$$\frac{y_0 + \Delta y}{y_0} = \frac{z_0 + \Delta z}{z_0} = \left(\frac{x_0 + \Delta x}{x_0}\right)^{-\nu_n}. \tag{S11}$$

Writing the deformation of the nucleus in the $x$-direction induced by the combination of pulling and pushing forces as $\Delta x_{\text{forces}}$, such that the new length of the nucleus is $x_{\text{forces}} = x_0 + \Delta x_{\text{forces}}$, we can then use Eq. (S11) to write the corresponding nuclear dimensions in the orthogonal directions as

$$y_{\text{forces}} = y_0 + \Delta y_{\text{forces}} = y_0 \left(1 + \frac{\Delta x_{\text{forces}}}{x_0}\right)^{-\nu_n} \tag{S12}$$

and

$$z_{\text{forces}} = z_0 + \Delta z_{\text{forces}} = z_0 \left(1 + \frac{\Delta x_{\text{forces}}}{x_0}\right)^{-\nu_n}, \tag{S13}$$

where $x_0/y_0/z_0$ denote the force-free width of the nucleus in the $x$ /$y$ /$z$-direction and $\nu_n$ is the Poisson ratio of the nucleus.

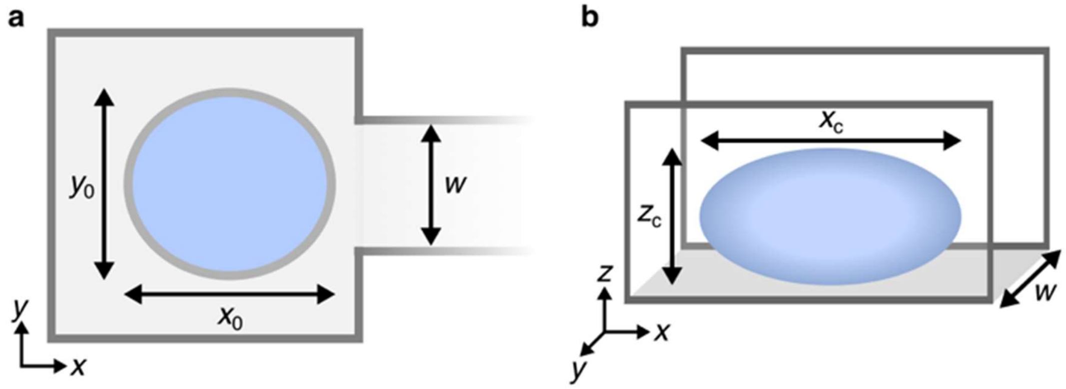


**Figure S6: Nuclear dimensions on the island (a) and in the channel (b).**

Now we add the confinement due to the channel of width $w$, such that the width of the nucleus in the channel $y_c = w$. Consequently, the diameter of the nucleus in the other dimensions is given by

$$x_c = x_{\text{forces}} \left(\frac{w}{y_{\text{forces}}}\right)^{-\nu_n} \tag{S14}$$

and

$$z_c = z_{\text{forces}} \left( \frac{w}{y_{\text{forces}}} \right)^{-\nu_n}. \tag{S15}$$

The aspect ratio between the two unconfined dimensions of the nucleus in the channel is then given by

$$AR_{\text{confined}} = \frac{x_c}{z_c} = \frac{x_0}{z_0} \left( \frac{x_{\text{forces}}}{x_0} \right)^{1+\nu_n}. \tag{S16}$$

If the pulling force is stronger than the pushing force, we expect that $\Delta x_{\text{forces}} > 0$ and thus $AR_{\text{confined}} > x_0/z_0 = AR_{\text{free}}$, while in the case that pushing forces are stronger than pulling forces, we expect $\Delta x_{\text{forces}} < 0$ and thus $AR_{\text{confined}} < AR_{\text{free}}$. We thus define the following shape parameter

$$\varepsilon = \frac{AR_{\text{confined}}}{AR_{\text{free}}} = \left( \frac{x_{\text{forces}}}{x_0} \right)^{1+\nu_n}. \tag{S17}$$

For $\varepsilon > 1$ pulling is the dominant force driving nucleus translocation, while for $\varepsilon < 1$ pushing dominates.

We do, however, not have direct experimental access to $x_{\text{forces}}$. To express $\varepsilon$ in terms of measurable quantities, we use that $x_{\text{forces}} = x_c \left( \frac{w}{y_{\text{forces}}} \right)^{\nu_n}$ and $y_{\text{forces}} = y_0 \left( \frac{x_{\text{forces}}}{x_0} \right)^{-\nu_n}$, such that

$$\frac{x_{\text{forces}}}{x_0} = \left[ \frac{x_c}{x_0} \left( \frac{w}{y_0} \right)^{\nu_n} \right]^{\frac{1}{1-\nu_n^2}} \tag{S18}$$

and thus

$$\varepsilon = \left[ \frac{x_c}{x_0} \left( \frac{w}{y_0} \right)^{\nu_n} \right]^{\frac{1+\nu_n^2}{1-\nu_n^2}}. \tag{S19}$$

#### 2.3. Deducing the force balance on the nucleus

To relate the observed nuclear shapes to the difference of pulling and pushing forces, we consider the force balance acting on the nucleus as it moves through the cytosol at constant speed (see Fig. S7). Since the drag force $F_{\text{drag}}$ acts on the center of mass of the nucleus, we split it up with $1/2F_{\text{drag}}$ acting on either side of the nucleus, such that overall the nucleus experiences a total drag force $F_{\text{drag}}$.

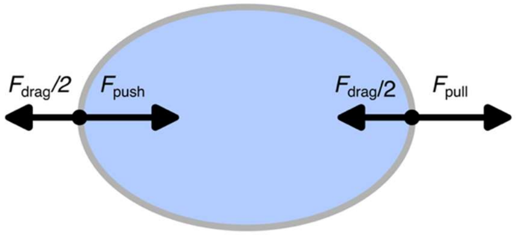


**Figure S7: Force balance on the nucleus.** The drag force is split over the front and the half of the nucleus, such that the overall drag force is applied to the centre of the nucleus.

We then get the following net force acting on the front and the back of the nucleus:

$$F_{\text{front}} = F_{\text{pull}} - \frac{1}{2}F_{\text{drag}} \tag{S20}$$

and

$$F_{\text{back}} = F_{\text{push}} - \frac{1}{2}F_{\text{drag}}. \tag{S21}$$

At a constant velocity and shape of the nucleus, the forces at the front and the back must balance out and be equal to the elastic force $F_{\text{deform}}$ generated by the nucleus due to its deformation:

$$F_{\text{deform}} = F_{\text{front}} = -F_{\text{back}} \tag{S22}$$

We can then rewrite $F_{\text{deform}} = (F_{\text{front}} - F_{\text{back}})/2$ in terms of the pulling and pushing forces as

$$F_{\text{deform}} = \frac{1}{2}\left(F_{\text{pull}} - F_{\text{push}}\right). \tag{S23}$$

As discussed above, the deformations of the nucleus can be approximated by the Hertz model, which yields the following expression of the deformation force in terms of the nuclear deformation along the direction of migration:

$$F_{\text{deform}} = \text{sign}\left(\Delta x^{\text{forces}}\right)\frac{4}{3}\frac{E_n}{1-\nu_n^2}\sqrt{\frac{x_0}{2}}\left|\frac{\Delta x^{\text{forces}}}{2}\right|^{3/2}. \tag{S24}$$

Together with Eq. (S18) the deformation force can then be written in terms of $\varepsilon$ as

$$F_{\text{deform}} = \text{sign}\left(\varepsilon^{1/(1+\nu_n)} - 1\right)\frac{1}{3}\frac{E_n}{1-\nu_n^2}{x_0}^2\left|\varepsilon^{1/(1+\nu_n)} - 1\right|^{3/2}. \tag{S25}$$

This expression allows us to estimate the nuclear deformation forces from the experimentally observed nuclear shapes.

#### 2.4. Plastic deformations and shape recovery

To analyse the viscoelastic response of nuclei getting deformed, we quantify the nuclear shapes depending on the channel width. We find that the shapes before and after the transition appear to be identical (Fig. S8a) and that the nuclei recover their shape within minutes after the transition (Fig. S8b). Unconfined nuclei appear to be slightly more elongated after transition through narrow channels (Fig. S8a).

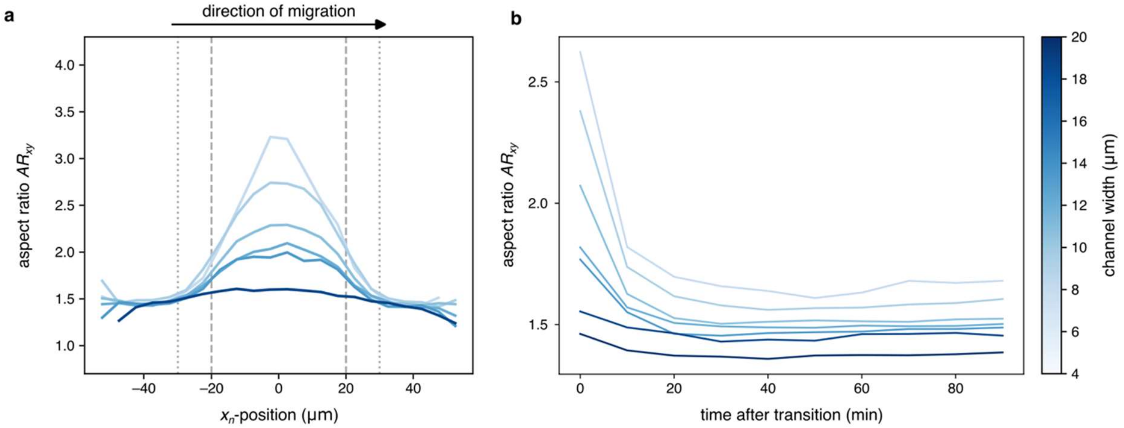


**Figure S8: Aspect ratio of the nucleus for different channel widths (x-dimension/y-dimension). (a)** Aspect ratio with respect to the nuclear position. The cells are oriented such that they migrate from left to right. The grey dashed lines indicate the edges of the channel and the grey dotted lines the point at which the nucleus starts entering (left)/ fully left (right) the channel. **(b)** Nuclear shape recovery after a transition.

### 3. Inference of the dynamical signature of the confinement

To obtain information about the effect that physical confinement has on the effective dynamics of migrating cells, we employ an approach that was successfully used to infer dynamical models from trajectories of cells migrating on two dimensional micro-patterns[8,9] and apply it to our case of cells migrating in 3D confinement. At the core of this inference approach lies the observation that the dynamics of migrating cells, quantified through the position of the nucleus along the long axis of the pattern $x_n$, can be effectively described by an underdamped Langevin equation[8]:

$$\ddot{x}_n = F(x_n, v_n) + \sigma\eta(t), \tag{S26}$$

Where $F(x_n, v_n)$ is the deterministic (drift) contribution to the nuclear acceleration and $\eta(t)$ is a gaussian white-noise random variable that accounts for random fluctuations in the nuclear dynamics. The strength of these random fluctuations is given by $\sigma$.

In particularly, we are interested in how the deterministic contribution $F(x_n, v_n)$ changes with decreasing channel width. The chosen experimental procedure produces micro-cavities with a continuous range of different channel widths. To have enough trajectories to perform the inference procedure at different channel widths, we bin the data according to the measured width of the pattern in the centre of the channel. The resulting number of trajectories for every bin together with the average bridge widths are summarized in Table S1.

To obtain a bias free estimate of the deterministic term $F(x_n, v_n)$, we employ a previously developed inference scheme, called Underdamped Langevin Inference[10]. This method is based on an expansion of $F(x_n, v_n)$ in terms of manually chosen basis functions $\hat{c}_{\alpha\beta}(x_n, v_n)$. The order to which this expansion is performed determines the number of parameters that have to be estimated from the data. This means that this method can produce many different estimators for $F(x_n, v_n)$, which requires selection criteria that ensure that the chosen model is sufficiently precise and unique for each bridge width.

#### 3.1. Symmetries of the basis functions

To reduce the number of free parameters as much as possible without constraining our self to a few leading order terms in the expansion of $F(x_n, v_n)$, we utilize the symmetries in our experimental setup to eliminate basis functions that do not match the systems symmetries. When choosing the centre of the pattern as the origin of our coordinate system, the symmetry of the pattern and the absence of any other anisotropic external cues implies that $F(-x_n, -v_n) = -F(x_n, v_n)$. To enforce this, we choose polynomial basis functions ${x_n}^l {v_n}^m$, where the combined order $l + m$ has to be odd.

#### 3.2. Criteria for the model selection

Even after the selection of a polynomial basis with the aforementioned symmetries, there is still freedom in the choice of the maximal included orders $l_{\max}$ and $m_{\max}$. Choosing $l_{\max}$ and $m_{\max}$ too low results in models of low complexity that might not be sufficient to capture the complex migration behavior. On the other hand, choosing $l_{\max}$ and $m_{\max}$ too high can lead to a number of model parameters that cannot be fully constrained from the data also resulting in a poor model. To select the right values for $l_{\max}$ and $m_{\max}$, we assess the predictive power of the inferred models.

The models are constrained by only using the instantaneous position, velocity and acceleration of the nucleus, without any knowledge about the dynamics on longer timescales. We use this to test our models by repeatedly simulating the inferred Langevin equation with random initial conditions to generate a synthetic dataset containing the same number of cells as the experimental dataset. When cells exit the experimentally sampled region in phase space we terminate the simulation for that cell. We then calculate a number of key statistics that describe the long timescale behaviour of the population of cells. These include the probability distribution of the nuclear position $p(x_n)$ and velocity $p(v_n)$ as well as the dwell time distribution $p(\tau)$.

To denoise the distributions, we estimate $p(x_n)$ and $p(v_n)$ in parameter-free way using the kernel-density estimate with Gaussian kernels implemented with the python library scipy[11], which closely matches the distributions obtained from histograms (see Fig. S9a, b). To estimate the dwell time distribution $p(\tau)$ one needs to account for the finiteness of the trajectories. We did this by using the python library livelines[12]. We found that a generalized gamma model was in close agreement with the parameter-free Kaplan-Maier estimator but was numerically more well behaved (see Fig. S9c), such that we chose this model for the calculation of $p(\tau)$.

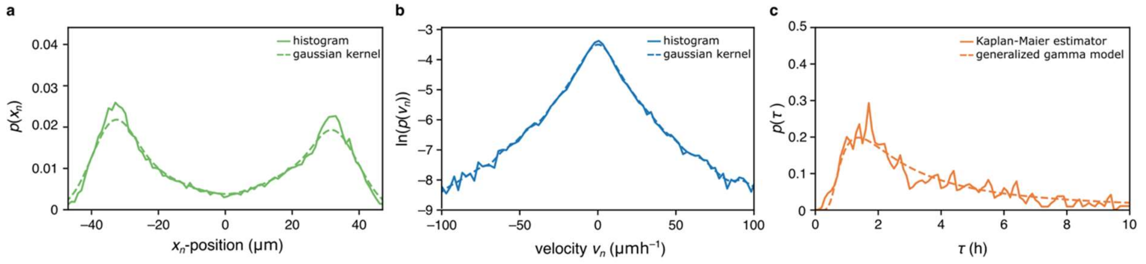


**Figure S9: Denoising of the experimental statistics used for model selection**. Considered were the probability distributions of (a) the nuclear velocities (b) the nuclear position (c) the dwell times.

The fact that all three of those quantities are probability distributions allows for a simple and comparable quantification of the difference between the experimental and the synthetic dataset. Due to its numerical stability and its boundedness, we choose to quantify the difference between the distributions obtained from the experiment and the simulations with the so-called Hellinger distance $H(p_{\exp}, p_{\inf})$[13], which takes values between 0 (perfect agreement between $p_{\exp}$ and $p_{\inf}$) and 1 (complete disagreement between $p_{\exp}$ and $p_{\inf}$). To quantify the accuracy of the inferred models, we sweep over $l_{\max}$ and $m_{\max}$ and infer models from one half of the available data (Fig. S10). We then simulate the model and calculate $p(x_n)$, $p(v_n)$ and $p(\tau)$ from this synthetic data set. We do the same for the half of the data that was not used for training and compute $H(p_{\mathrm{exp}}, p_{\mathrm{inf}})$ for all three statistics. As shown in Fig. S10, there are strong differences in the performance of the different models.

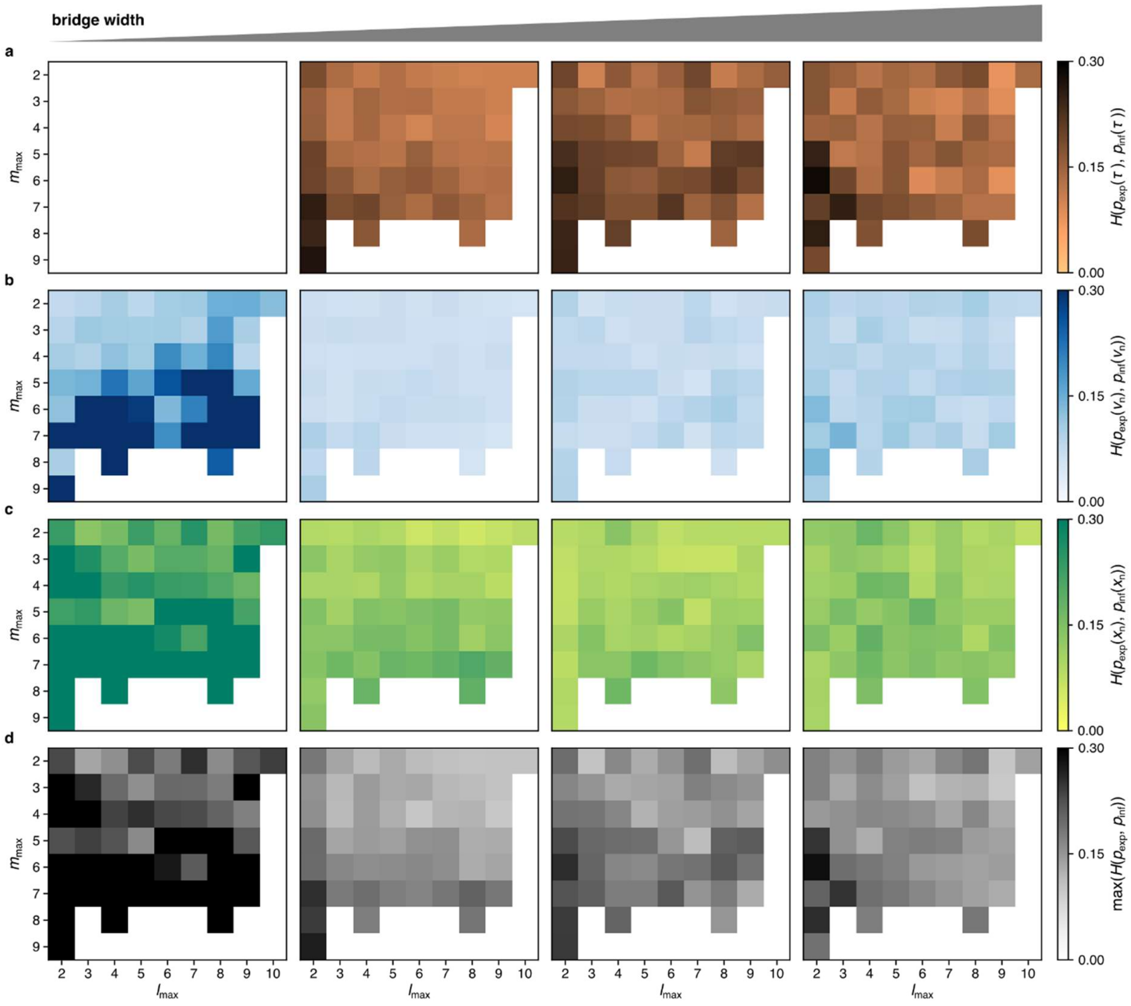


**Figure S10: Overview of the Hellinger distances between experimental and model statistics for models of different orders. (a)** Hellinger distances of the dwell time distributions. At the narrowest channel width, transitions were so rare that a dwell time distribution could not be computed reliably. **(b)** Hellinger distances of the velocity distributions. **(c)** Hellinger distances of the position distributions. **(d)** Maximal Hellinger distance of the three considered statistics.

#### 3.3. Uniqueness of the inferred models

To choose a value for $H_{\mathrm{thresh}}$, we started out with a value of $H_{\mathrm{thresh}} = 0.15$, which ensures that already large parts of the probability distributions are reproduced by the model. A model was accepted if all three probability distributions scored a value of $H(p_{\mathrm{exp}}, p_{\mathrm{inf}}) < H_{\mathrm{thresh}}$. We found however that this criterium was not strict enough to lead to unique models, but instead found qualitative differences between the different models. We then gradually decreased the threshold until we obtained a selection of quantitively identical models for each bridge width. This was the case for a value of $H_{\mathrm{thresh}} = 0.13$ (see Fig. S11 for $H_{\mathrm{thresh}} = 0.14$ and Fig. S12 for $H_{\mathrm{thresh}} = 0.13$).

For the narrowest bridge widths ($w \approx 5$ µm) almost no transitions were observed anymore. As a consequence, the dwell time distribution could not be properly estimated and was excluded from as a selection criterium for this bridge width. Further, the model was not able to reproduce the sharply peaked positional probability distributions well enough to pass the threshold. Since we are however more interested in the dynamical properties of our model, we loosened the selection criterium in that case to $H_{thres} \ = 0.14$. The models that were shown in the main text are given in Table S2.

| Bridge width | 4 µm | 7 µm | 9 µm | 12 µm |
|---|---|---|---|---|
| $l_{\max}$ | 3 | 9 | 8 | 7 |
| $m_{\max}$ | 2 | 3 | 2 | 3 |

**Table S2: Orders of the basis functions used to infer the models discussed in the main text.**

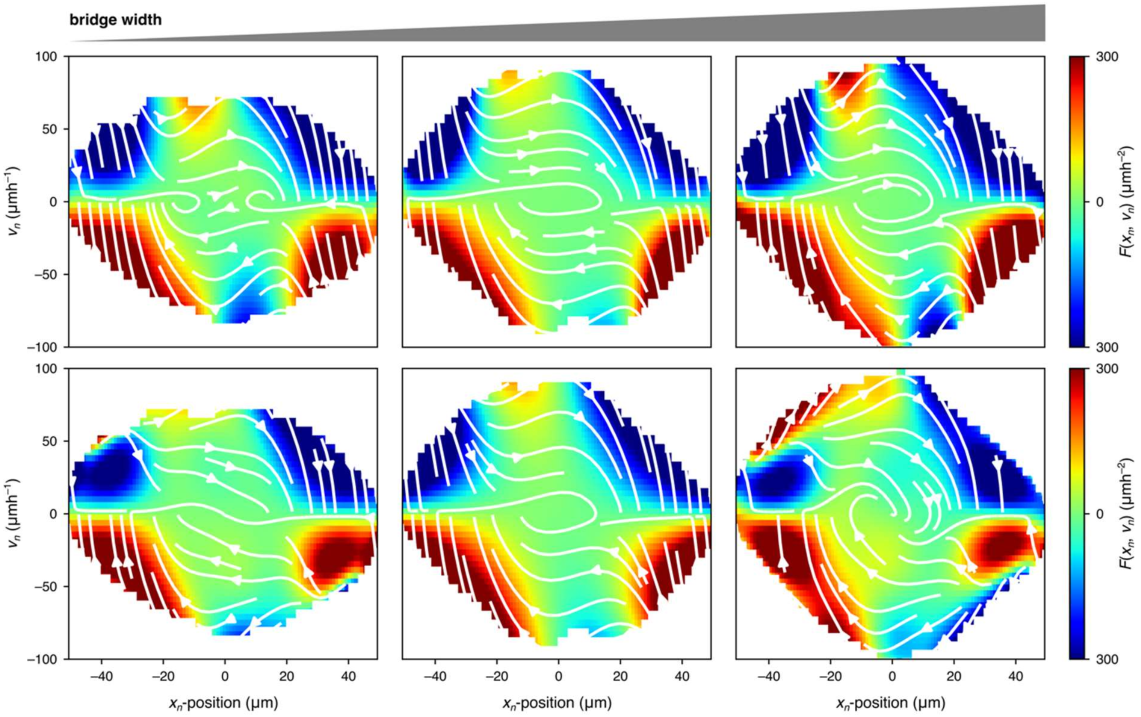


**Figure S11: Comparison between exemplary models with a threshold of 0.14.** Shown are the bridge widths 7 µm, 9 µm and 12 µm.

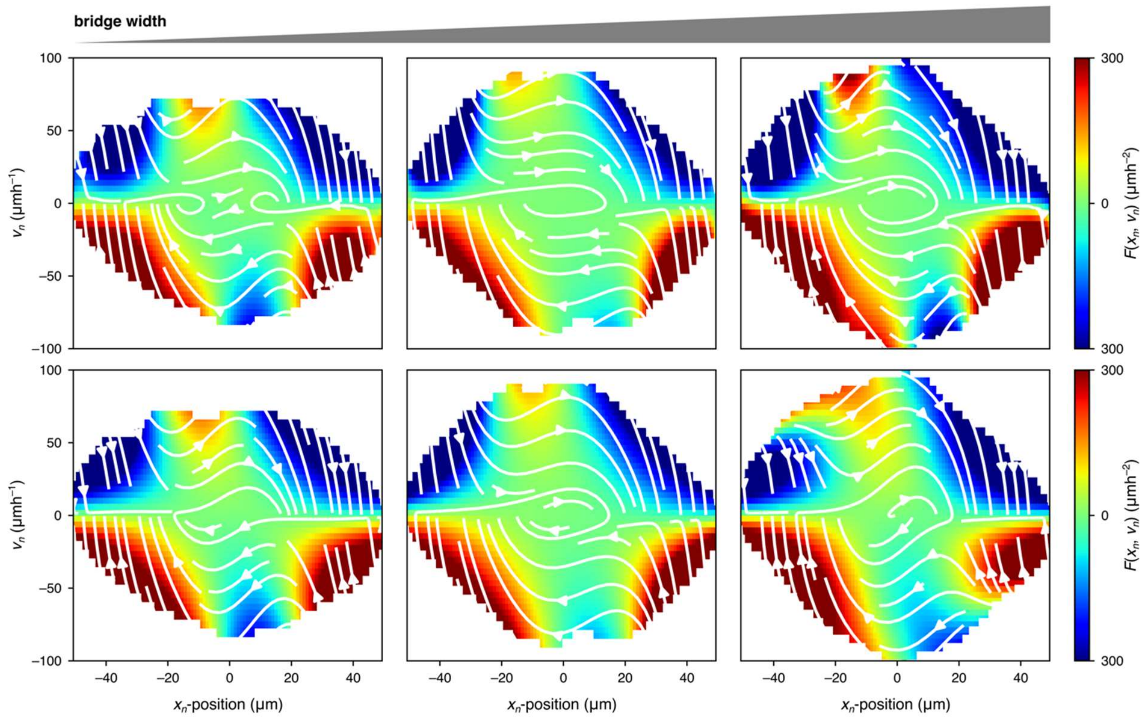


**Figure S12: Comparison between exemplary models with a threshold of 0.13.** Shown are the bridge widths 7 µm, 9 µm and 12 µm.

### 4. Mechanistic model

To model the effect of 3D confinement on mesenchymal cell migration, we adapt a model for cell migration on 2D dumbbell-shaped micro-patterns that we previously constrained from experimental data[9].

#### 4.1. General structure of the model

The model accounts for three central degrees of freedom of the cell: the nuclear position, the position of the leading protrusion of the cell and the polarity of the cell that drives protrusion growth (see Fig. 5c in the main text). The coupling between nucleus and protrusion is modelled by linear elastic spring connecting the two. The dynamics of the nucleus are then given by the following equation of motion:

$$\gamma(x_n)\dot{x}_n = k_n(x_p - x_n) + f_{\text{push}}(x_n) + f_{\text{channel}}(x_n) \tag{S27}$$

Here, $\gamma(x_n)$ accounts for spatial variations in the nuclear friction coefficient due to reduced amount of adhesion bonds in the channel compared to the islands, $k_n$ denotes the effective spring constant that acts on the nucleus, $f_{\text{push}}(x_n)$ is the pushing force generated by rear contractions of the cell and $f_{\text{channel}}(x_n)$ denotes the force due to the nuclear deformations induced by the channel.

The dynamics of the protrusion coordinate are given by

$$\dot{x}_p = -k_p(x_p - x_n) - \partial_{x_p} V(x_p) + rP(t), \tag{S28}$$

where $k_p$ is the effective spring constant that acts on the nucleus. The protrusion formation is limited to the region of the dumbbell-shaped micro-pattern. To account for this, a repulsive potential $V(x_p) = (x_p/x_{\text{boundary}})^8$ is introduced that acts on the protrusion coordinate. Additionally, to account for the internal organization of the cell, a polarization $P(t)$ acts on the protrusion coordinate driving protrusion formation in the direction of polarization. The cell polarity $P$ is converted into a protrusion growth rate through the parameter $r$.

In previous work, we found that the polarization is sensitive to the geometry of the pattern[9]: In the absence of confinement, the cell is on average unpolarized and $P(t)$ fluctuates around 0. In contrast, in the case of narrow confinement of the protrusion, the cell polarization gets reinforced and $P(t)$ takes a finite value. This behavior is summarized in the following equation:

$$\dot{P} = -\alpha(x_p)P - \beta P^3 + \sigma\xi(t) \tag{S29}$$

Here, the geometry sensitive parameter determines the value of the stable fixed point $P^*$ of $P(t)$ with $P^* = \pm\sqrt{-\alpha/\beta}$ for $\alpha < 0$ and $P^* = 0$ for $\alpha \geq 0$. The constant $\beta > 0$ ensures that $P(t)$ remains bounded at all times. The term $\sigma\xi(t)$ denotes a Gaussian white noise-process of strength $\sigma$ and with $\langle\xi(t)\rangle = 0$ and $\langle\xi(t)\xi(t')\rangle = \delta(t - t')$ that accounts for stochastic fluctuations in the internal organization of the cell.

To account for the increased confinement in the centre of the pattern, we gradually decrease $\alpha$ towards the center of the pattern according to

$$\alpha(x_p) = \begin{cases} \alpha^{\min} + \frac{\alpha_0 - \alpha^{\min}}{2}\left(1 - \cos\left(\frac{x_p \pi}{L_\alpha}\right)\right), |x_p| < L_\alpha \\ \alpha_0, \quad \text{else} \end{cases}. \tag{S30}$$

The parameter $\alpha^{\min}$ decreases with decreasing bridge width according to[14]

$$\alpha^{\min} = -\alpha_1 + \alpha_2 w. \tag{S31}$$

For channel widths beyond the unconfined width of the cell $w_{\text{free}}$, we expect the channel width to only have a minimal effect on the experiment, such that in that regime, we assume that $\alpha^{\min} = \alpha_0$.

#### 4.2. Nuclear friction

In the absence of nuclear deformations, the nuclear friction was found to be reduced in the centre of dumbbell shaped patterns[9]. A possible explanation for this is the reduced number of focal adhesions due to the reduced adhesive area in the constriction[14]. In the presence of physical walls that strongly confine the nucleus, however, we expect that the nuclear friction increases in the channel due to interactions with the walls. To account for the varying friction that the nucleus experiences on the pattern, we gradually adapt $\gamma(x_n)$ from its value of 1 on the island towards a value $\gamma^{\text{centre}}$ in the centre of the pattern. For this we follow the approach used in Brückner et al.[9] and use the following analytic expression for $\gamma(x_n)$:

$$\gamma(x_n) = \gamma^{\text{centre}} + \frac{1 - \gamma^{\text{centre}}}{2}\left(1 - \cos\left(\frac{x_n \pi}{L_{\text{sys}}}\right)\right) \tag{S32}$$

Here, $2L_{\text{sys}}$ denotes the total length of the pattern.

If the cell is completely unconfined ($w \geq w_{\text{free}}$), we simply expect $\gamma^{\text{centre}} = 1$.

When the cell is confined but the nucleus is not deformed ($2R_n \leq w < w_{\text{free}}$), we expect a linear decrease of $\gamma^{\text{centre}}$ with channel width due to the reduced adhesive area compared to the unconfined case[9,14]. Finally, in the presence of nuclear deformations ($w < 2R_n$)), we expect a non-linear increase of the nuclear drag with reducing channel width[15]. Taken together, we use the following expression for the nuclear friction in the channel:

$$\gamma^{\text{centre}} = \begin{cases} 1, w > w_{\text{free}} \\ \frac{1 + \gamma_1 w}{1 + \gamma_1 w_{\text{free}}}, 2R_n < w \leq w_{\text{free}} \\ \frac{1 + 2\gamma_1 R_n}{1 + \gamma_1 w_{\text{free}}}\left(1 + \frac{\gamma_2}{w}\right), 2R_n \geq w \end{cases} \tag{S33}$$

#### 4.3. Deformation barrier

Motivated by the observation that the 3D channel appears as an effective barrier in the inferred dynamics of the nucleus for channel widths below the size of the nucleus (see Fig. 3 in the main text) together with the steep decrease in transition rates in that regime, we account for nuclear deformations induced by the channel through a potential barrier, such that

$$f_{\text{channel}}(x_n) = -\partial_{x_n} W(x_n). \tag{S34}$$

As the nucleus moves into the confinement, it is compressed in the orthogonal direction, which leads to a lengthening of the nucleus. The confinement exerts a force opposite to the direction of movement onto the nucleus until it has completely entered the bridge. We thus use the following expression for the potential barrier:

$$W(x) = W_{\max} \begin{cases} \frac{x}{L_W} - \pi^{-1} \cos\left(\frac{\pi x}{L_W}\right) \sin\left(\frac{\pi x}{L_W}\right), 0 < x < L_W \\ 0, \qquad\qquad\qquad\qquad\qquad \text{else} \end{cases}, \tag{S35}$$

where $L_W = L_b/2 + R_n$ with $R_n$ being the average radius of the unconfined nucleus. Since the pattern and consequently also the potential is symmetric, we can calculate the value of the potential barrier at position $x_n$ through $W(x = L_W - |x_n|)$. We thus get the channel-induced deformation force

$$f_{\text{channel}}(x_n) = -\partial_x W(x) \partial_{x_n} x|_{x=L_W-|x_n|} = \text{sign}(x_n) \partial_x W(x)|_{x=L_W-|x_n|}. \tag{S36}$$

The observed 3/2-scaling of the nuclear normal forces with the nuclear deformations represents a lower bound for the dependence of the force opposing the nuclear deformations. We thus used the following expression for $W_{\max}$, which yielded good agreement between our model and the experiments:

$$W_{\max} = W_0 (2R_n - w)^{3/2} \tag{S37}$$

#### 4.4. Pushing force

There is recent evidence that in confinement, cells can generate pushing forces to move the nucleus through a constriction[16,17]. In the absence of confinement, we assume that the pushing force can be neglected due to the unhindered fluid exchange between the front and the rear of the cell. To account for the gradual increase in nuclear deformation, we apply an increasing pushing force that acts directly onto the nucleus as the nucleus moves into the centre of the channel. The pushing force is then given by

$$f_{\text{push}}(x_n) = \text{sign}(x_p - x_n) f_{\max}(w) \cos\left(\frac{\pi x_n}{L_b}\right). \tag{S38}$$

To find an expression for $f_{\max}(w)$, we consider the case of a nucleus that is completely confined to a width $w$ in the $y$-direction. These pushing forces are thought to be triggered by strong nuclear deformations that lead to $Ca^{2+}$ release and consequently increased myosin recruitment to the cortex[18,19]. This leads to increased cortical tension and thus an increased pushing force due to the increased Laplace pressure. The pushing force in confinement is then proportional to the increase in cortical tension $\Delta\tau_{\mathrm{cortex}}$ and the projected area $S$ of the nucleus in the plane orthogonal to the direction of migration[20]

$$f_{\max}(w) \propto S\big(w(x_n)\big)\Delta\tau_{cortex}. \tag{S39}$$

Since we expect the dominant contribution to the bridge width dependence of $S\big(w(x_n)\big)$ to be the compression imposed by the side walls of the channel, we can estimate the projected area of the nucleus as

$$S\big(w(x_n)\big) = S_0\left(\frac{w(x_n)}{2R_n}\right)^{1-\nu_n}, \tag{S40}$$

where $S_0$ is the projected area of the undeformed nucleus.

The calcium release upon physical confinement of the nucleus is associated with stretch-sensitive calcium channels in the perinuclear endoplasmic reticulum[19]. To relate the change in $Ca^{2+}$ concentration in the cytosol to the degree of confinement, we assume that the concentration is proportional to the probability that a channel is open. We use a simple mechanical model for the opening probability of the calcium channels, which gives us[21]

$$n^{Ca^{2+}} \propto \frac{1}{1+\exp((\Delta E - \Delta A\tau_{ER}(w(x_n))/k_B T)}, \tag{S41}$$

where $\Delta E$ is the energy difference between the opened and closed state of the calcium channel and $\Delta A$ is the change in area of the channel upon opening, caused by the tension in the perinuclear endoplasmic reticulum $\tau_{ER}$.

We assume a linear scaling of $\tau_{ER}$ with the nuclear deformation $2R_n - w(x_n)$ to leading order. In the range of nuclear deformations probed in our experiments, we expect the exponential part of the sigmoidal function in the previous equation dominates. This assumption is supported by measurements of the cortical myosin concentration under confinement in primary progenitor stem cells cultured from zebrafish embryos[18], which probed confinement heights down to 7 µm without observing a clear deviation from an exponential dependence. We can then write in $Ca^{2+}$ concentration as

$$\Delta n^{Ca^{2+}} = n_0^{Ca^{2+}}\left(e^{\frac{2R_n - w(x_n)}{h^*}} - 1\right), \tag{S42}$$

where $h^*$ is the characteristic confinement height of calcium release.

In principle, the cortical tension could show a non-linear response on the calcium concentration due to effects like contraction induced Rho release[16] or saturation of the contractility. However, to keep our model simple, we only consider a linear dependence of the cortex tension on the calcium concentration. We checked that allowing for higher order terms does not change our results qualitatively. We can then express the pushing force explicitly as a function of the confinement width of the nucleus $w(x_n)$ as

$$f_{max}(w) = S_0 \tau^0_{\text{cortex}} \left(\frac{w(x_n)}{2R_n}\right)^{1-\nu_n} \left(e^{\frac{2R_n - w(x_n)}{h^*}} - 1\right)$$
$$\approx S_0 \tau^0_{\text{cortex}} \left(\frac{w(x_n)}{2R_n}\right)^{1-\nu_n} \frac{2R_n - w(x_n)}{h^*} \quad \text{(S43)}$$

where $\tau^0_{cortex}$ denotes the cortical tension in the absence of nuclear deformations.

#### 4.5. Implementation and simulations

The set of closed Eqs. (27) – (29) was implemented in Python and numerically solved with a standard forward Euler scheme. To ensure comparability to the experimental data, we sampled values every 10 min for the simulated trajectories and chose the trajectory length such that it corresponded to 50 h, which is similar to the maximum observation time in the experiment.

Whenever possible the values of the model parameters were taken previous work[9,14] or set to match the experimental geometry. The remaining parameters used in the simulations were constrained by fitting the model to a number of key experimental statistics (nuclear deformation forces, transition rates and nuclear velocities). The resulting parameter values are given in Table S4.

| Parameter | value | parameter | value |
|---|---|---|---|
| $L_{\text{sys}}$ (μm) | 55 (experiment) | $R_n$ (μm) | 6 (experiment) |
| $L_c$ (μm) | 40 (experiment) | $k_n$ ($\text{h}^{-1}$) | 1.2 (fit) |
| $x_{\text{boundary}}$ (μm) | 0.4 × $L_{sys}$ [9]) | $k_p$ ($\text{h}^{-1}$) | 2.4 (fit) |
| $\sigma$ ($\mu\text{mh}^{-3/2}$) | 100 [9] | $L_\alpha$ (μm) | 25 (fit) |
| $\alpha_0$ ($\text{h}^{-1}$) | 1.1 (fit) | $W_0$ ($\mu\text{m}^{1/2}\ \text{h}^{-1}$) | 40 (fit) |
| $\alpha_1$ ($\text{h}^{-1}$) | 2.1 (fit) | $\frac{S_0 \tau^0_{cortex}}{h^*}$ ($\mu\text{m h}^{-1}$) | 16 (fit) |
| $\alpha_2$ ($\text{h}^{-1}$) | 0.005 (fit) | $r$ | 2 (fit) |
| $\beta$ ($\text{h}^{-1}$) | 0.0001 [9] | $\gamma_1$ ($\mu\text{m}^{-1}$) | 0.3 [14] |
| $w_{\text{free}}$ (μm) | 25 (experiment) | $\gamma_2$ ($\mu\text{m}^{-1}$) | 15 (fit) |

**Table S3: Model parameters.**

#### 4.6. Calculation of the effective underdamped dynamics

To compare the simulated dynamics with the inferred dynamics, we need to find a formulation of the model in terms of an underdamped Langevin equation (Eq. (1) in the main text). To derive an expression of the deterministic term $F(x_n, v_n)$ from our mechanistic model, we follow previous work[9] and rewrite the equations for the nuclear and the protrusion coordinate as

$$\gamma(x_n)\dot{x}_n = k_n\big(x_p - x_n\big) + f_{\text{push}}(x_n) + f_{\text{channel}}(x_n) \equiv f_n(x_n, x_p) \tag{S44}$$

and

$$\dot{x}_p = -k_p\big(x_p - x_n\big) - \partial_{x_p}V\big(x_p\big) + rP(t)) \equiv f_p(x_n, x_p) + rP(t). \tag{S45}$$

This allows us to calculate $F(x_n, v_n)$ according to Eq. (1) from the main text as

$$\begin{aligned} F(x_n, v_n) = \langle \dot{v}_n | x_n, v_n \rangle &\overset{\text{Eq. (S44)}}{=} \left\langle \frac{df_n\big(x_n, x_p\big)}{dt} \middle| x_n, v_n \right\rangle = \left\langle \frac{df_n}{dx_n} v_n + \frac{df_n}{dx_p} \dot{x}_p \middle| x_n, v_n \right\rangle \\ &\overset{\text{Eq. (S45)}}{=} v_n \frac{df_n}{dx_n} + \frac{df_n}{dx_p} \big\langle f_p(x_n, x_p) + rP(t) \big| x_n, v_n \big\rangle \\ &\overset{\text{Eq. (S44)}}{=} v_n \frac{df_n}{dx_n} + \frac{df_n}{dx_p} f_p(x_n, x_p)\Big|_{x_p = x_n + k_n^{-1}(\gamma(x_n)v_n - f_{\text{push}}(x_n) - f_{\text{channel}}(x_n))} \\ &\quad + \frac{df_n}{dx_p} \langle rP(t) | x_n, v_n \rangle \end{aligned} \tag{S46}$$

While most terms in Eq. (S46) can be calculated analytically, $\langle rP(t) | x_n, v_n \rangle$ needs to be determined numerically. The resulting $F(x_n, v_n)$ and corresponding confinement signature maps are shown in Fig. S23b and c.

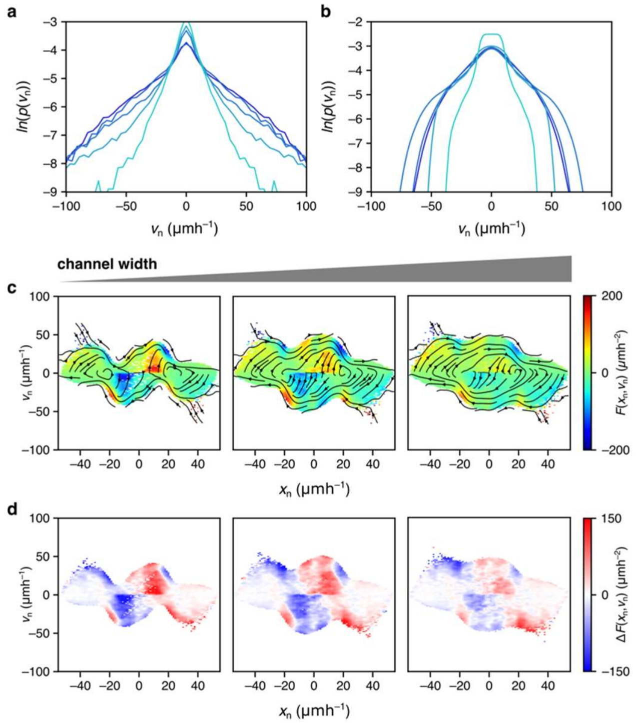


**Figure S13: Further comparison between simulation and experiment. (a), (b)** Probability distributions of the nuclear velocities at varying channel widths obtained experimentally (a) and via simulations (b) (same colours as Fig. 2a,b in the main text). **(c), (d)** Effective underdamped dynamics of the model. Deterministic contribution (c) and corresponding confinement signature maps (d) of the effective underdamped dynamics of the model (from left to right: 4 µm, 7 µm, 9 µm).

#### 4.7. Comparison of experimental data and model predictions

We find that both the distribution of the nuclear position and velocity show semi-quantitative agreement with the experimental data. In particular, the model captures the overall dependence of the distributions on the channel width, with the appearance of clear peaks of the $x_n$-distribution in the chambers and pronounced minima in the channel at narrow channel widths (main text Fig. 2b). Further, we observe an overall narrowing of the nuclear velocities with increasing confinement (Fig. S13a,b). We find that the location of the peaks in the $x_n$-distribution deviates between experiment and simulations. For the simulation we find a maximum near the entrance of the channel, while in the experiments the maximum is located more towards the centre of the chamber. In line with this, we find that the biggest differences between the inferred dynamics and the effective underdamped dynamics of the model are found on the islands, while our model captures the dynamics of the actual transition accurately (main text Fig. 3). From assessment of the bright-field videos, we found that after a failed transition attempt during which the nucleus is placed near the entrance of the channel, the cells often retract their nucleus back to the centre of the channel for an extended period of time before it starts a new attempt. Since our model is designed to capture the interactions between the confinement and the cell and not necessarily the cellular behaviour in the chamber, we did not include a mechanism that could account for this in our model in order to keep our mechanistic model as simple as possible while still capturing the essence of the cell confinement interactions. Beyond the differences between the model and the experiments in the chambers, we also find that the experimental velocity distribution fall off slower (Fig. S23a,b). This does however mostly affect the region of extremely high velocities (>50 μm $h^{-1}$), which occur extremely rare, such that they have little effect on the population averaged migration dynamics.
